# Integrating single-cell RNA-seq datasets with substantial batch effects

**DOI:** 10.1101/2023.11.03.565463

**Authors:** Karin Hrovatin, Amir Ali Moinfar, Luke Zappia, Alejandro Tejada Lapuerta, Ben Lengerich, Manolis Kellis, Fabian J. Theis

## Abstract

Integration of single-cell RNA-sequencing (scRNA-seq) datasets has become a standard part of the analysis, with conditional variational autoencoders (cVAE) being among the most popular approaches. Increasingly, researchers are asking to map cells across challenging cases such as cross-organs, species, or organoids and primary tissue, as well as different scRNA-seq protocols, including single-cell and single-nuclei. Current computational methods struggle to harmonize datasets with such substantial differences, driven by technical or biological variation. Here, we propose to address these challenges for the popular cVAE-based approaches by introducing and comparing a series of regularization constraints.

The two commonly used strategies for increasing batch correction in cVAEs, that is Kullback–Leibler divergence (KL) regularization strength tuning and adversarial learning, suffer from substantial loss of biological information. Therefore, we adapt, implement, and assess alternative regularization strategies for cVAEs and investigate how they improve batch effect removal or better preserve biological variation, enabling us to propose an optimal cVAE-based integration strategy for complex systems. We show that using a VampPrior instead of the commonly used Gaussian prior not only improves the preservation of biological variation but also unexpectedly batch correction. Moreover, we show that our implementation of cycle-consistency loss leads to significantly better biological preservation than adversarial learning implemented in the previously proposed GLUE model. Additionally, we do not recommend relying only on the KL regularization strength tuning for increasing batch correction, as it removes both biological and batch information without discriminating between the two. Based on our findings, we propose a new model that combines VampPrior and cycle-consistency loss. We show that using it for datasets with substantial batch effects improves downstream interpretation of cell states and biological conditions. To ease the use of the newly proposed model, we make it available in the scvi-tools package as an external model named sysVI. Moreover, in the future, these regularization techniques could be added to other established cVAE-based models to improve the integration of datasets with substantial batch effects.

## Introduction

The joint analysis of multiple single-cell RNA sequencing (scRNA-seq) datasets has recently provided new insights that could not have been obtained from individual datasets. For example, pooling of datasets generated in different studies enabled cross-condition comparisons^1,2^, population-level analysis^3,4^, and revealed evolutionary relationships between individual cell types^5^. The selection of pre-clinical models, such as organoids and animals, and the characterization of their limitations likewise rely on comparison with human tissues^6–14^. Similarly, the selection of the optimal sequencing protocols requires a comparison of datasets generated with different protocols^15,16^. Lastly, currently arising large-scale “atlases” that are posed to serve as references of cell biology are aimed to combine public datasets with substantial technical and biological variation, including multiple organs and developmental stages^17^. Overall, with the increasing number of publicly available scRNA-seq datasets^18^, the number of such cross-dataset analyses is also increasing.

These analyses can be complicated due to technical and biological differences between samples^19–21^. To overcome this, computational methods for single-cell specific data integration have been developed^22,23^ and previous benchmarks have evaluated their integration performance generally^20^ and on cross-species datasets specifically^19^. Among the most popular and best-performing methods are conditional variational autoencoder (cVAE) based models, which are able to correct non-linear batch effects, are flexible in the choice of batch covariates, and are particularly scalable to large datasets^20,22^. Thus, they are also a method of choice for single-cell atlases^4,24,25^. However, while cVAE-based and other non-deep-learning methods enable good integration of batch effects caused by processing similar samples in different laboratories, they do not enable sufficient integration when differences between datasets are more substantial due to datasets originating from distinct biological or technical “systems”, such as multiple species or sequencing technologies (e.g. cell-nuclei)^19–21^ (**Figure 1a,b**), as demonstrated later. To enable more comprehensive single-cell atlasing efforts that will integrate diverse samples with stronger batch effects, it is thus vital to further improve the performance of the commonly used cVAE-based integration models.

**Figure 1:**
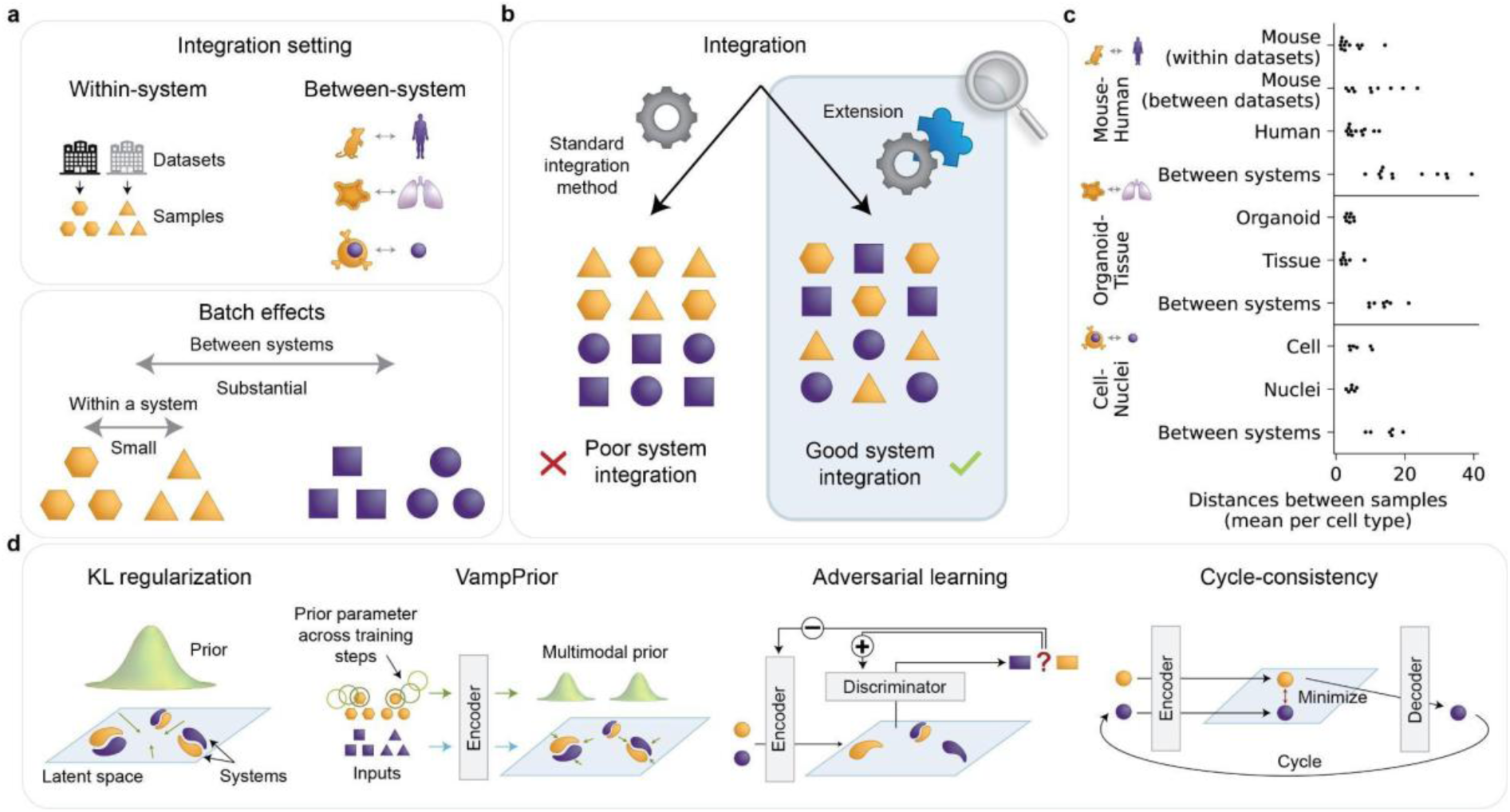
The challenge of integrating datasets with substantial batch effects. (**a**) Substantial batch effects are present between different biological “systems”, such as cross-species, organoid-tissue, and cell-nuclei datasets. We selected one dataset for each of these three between-system integration types as shown in panel **c**. (**b**) Integrating datasets with substantial batch effects poses a bigger challenge than integrating datasets of similar samples across laboratories, where the batch effect is smaller. In this study, we evaluate different approaches for improving cVAE-based batch correction. (**c**) Pre-integration distances between samples from the same or different systems. Points show mean per cell type. (**d**) Overview of approaches for improving batch removal in cVAE-based models: KL-loss-based regularization of the latent space, the use of the VampPrior as a replacement for the standard Gaussian prior, and adversarial learning and cycle-consistency loss that actively push together samples from different systems. Parts of the cVAE model were omitted from individual panels for brevity. For a detailed description of the standard cVAE and its extensions, see the methods section *Overview of cVAE-based integration approaches*.

Increased batch integration can be achieved via multiple extensions of cVAE models, including Kullback–Leibler divergence (KL) regularization strength tuning^26^, batch distributions alignment approaches^27–29^ (with latent space adversarial learning^30–32^ being the prime example), and latent space cycle-consistency that was previously only used for multi-omic integration in combination with adversarial learning^33–35^ (**Figure 1d**). For increasing biological preservation in scRNA-seq representation learning without supervision, we previously proposed the use of the multimodal variational mixture of posteriors (VampPrior)^36^ as the prior for the latent space (**Figure 1d**) in a workshop paper^37^, but this extension has not been explored in integration models. While cycle consistency and the VampPrior have been previously applied to single-cell data, it is unclear what their strengths and weaknesses are for integration and how they compare when used under different data scenarios.

Here, we explore the shortcomings of popular cVAE-based integration strategies, namely KL regularization tuning and adversarial learning, and show how these can be overcome by using cycle-consistency loss and the VampPrior. We systematically evaluate batch correction and biological preservation on cell type and sub-cell type levels, with existing^20^ metrics and a newly proposed metric for assessing within-cell-type variation. We study these across three scenarios with substantial batch effects: cross-species, organoid-tissue, and cell-nuclei. In short, we find that the combination of the VampPrior and cycle-consistency (VAMP+CYC model) improves batch correction while retaining high biological preservation, making VAMP+CYC the method of choice for integrating datasets with substantial batch effects. The VAMP+CYC model also empowers post-integration analysis of cell states and biological conditions. Our model is easily accessible to the community as part of the sciv-tools package^38^ under the name sysVI, short for “integration of diverse <u>sys</u>tems with <u>v</u>ariational inference”, and the here-proposed strategies could be likewise easily added to other cVAE-based integration methods.

## Results

We explored shortcomings of existing methods for integrating substantial batch effects and subsequently developed an improved method to suit diverse use cases. We selected multiple datasets where batch effects are more substantial than what is commonly observed between samples within a dataset or between similar datasets generated in different laboratories (Figure 1a**,c**). This includes three between-system data use cases: organoids (N samples = 21, N cells = 43,505) and adult human tissue samples (N samples = 20, N cells = 54,491) of the retina (organoid-tissue), scRNA-seq (N samples = 9, N cells = 28,465) and single-nuclei RNA-seq (snRNA-seq) (N samples = 9, N nuclei = 57,599) from subcutaneous adipose tissue (cell-nuclei), and mouse (N datasets = 8, N samples = 52, N cells = 263,140) and human (N samples = 65, N cells = 192,203) pancreatic islets (mouse-human). We confirmed that in all three data cases, the per-cell type distances between samples on non-integrated data are significantly smaller within systems, both within and between datasets, than between systems (Figure 1c, **Supplementary Table S1**).

## Existing methods struggle with the loss of biological information when increasing batch correction

Tuning of KL regularization strength is the most widely adopted approach for tuning batch correction strength as it is part of the standard cVAE architecture. It regulates how much the cell embeddings may deviate from the standard Gaussian distribution. By definition, KL regularization does not distinguish between biological and batch information, jointly removing both of them. To assess this, we measured batch correction via graph integration local inverse Simpson’s index (iLISI)^20^, which evaluates batch composition in the local neighborhoods of individual cells, and cell-type level biological preservation with a modified version of normalized mutual information (NMI)^20^ metric, which compares clusters from a single clustering resolution (NMI fixed) to grounded-truth annotation. Indeed, increased KL regularization strength led to higher batch correction and lower biological preservation (Figure 2a, **Supplementary Figure S1**). In particular, we observed that increased KL regularization strength led to some latent dimensions being set close to zero in all cells, resulting in information loss (**Supplementary Figure S2**). Thus, the higher batch correction score is simply a consequence of fewer embedding dimensions that were effectively used in downstream analyses. This was reflected by standard scaling of individual embedding features resulting in the loss of KL regularization-induced changes in integration scores in all three data use cases (Figure 2a, **Supplementary Figure S3**, **Supplementary Note S1**). In particular, on the scaled data higher KL regularization strength did not lead to increased batch correction or reduced biological preservation. Thus, KL weight is not a favorable approach for removing batch effects as it removes both biological and batch variation without discrimination.

**Figure 2:**
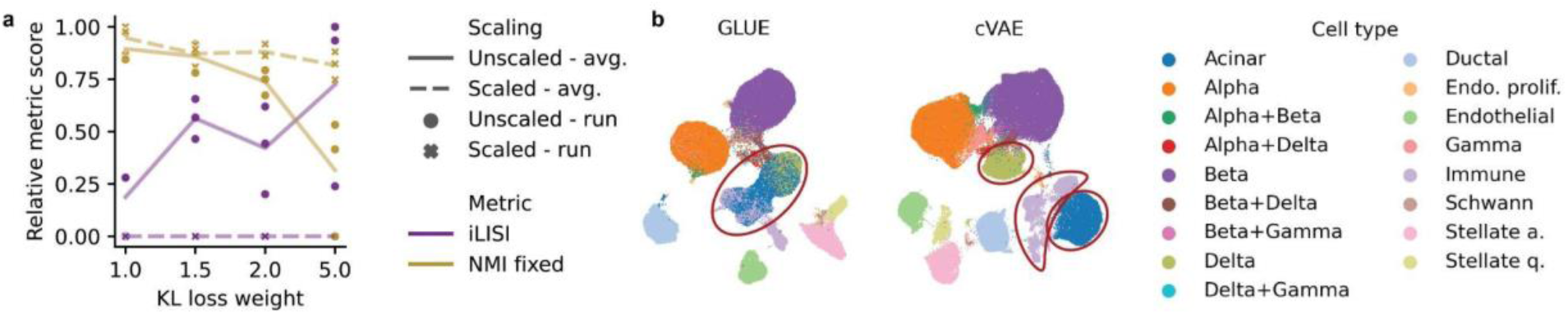
Adversarial learning and KL regularization strength tuning struggle with retaining biological variation when increasing batch correction. (**a**) Integration performance of scaled and unscaled cVAE embedding with different KL regularization loss weights. Shown are batch correction (iLISI, higher is better) and cell type level biological preservation (NMI fixed, higher is better) metrics scaled to [0,1] per metric. Individual runs with different seeds are shown as points and their averages (avg.) as lines. Results are presented for mouse-human data, with similar performance trends observed in other data use cases as reported in **Supplementary Figure S4**. (**b**) UMAP visualization of mouse-human dataset integrated with standard cVAE model and GLUE with the best hyperparameter setting, colored by cell type. Red circles mark examples of cell types that are mixed in the GLUE but not the cVAE integration.

The most popular cVAE extensions for actively pushing together cells from different batches are based on batch distribution alignment^27–29^, especially adversarial learning^30–32,39,40^. However, these approaches are prone to mix embeddings of unrelated cell types with unbalanced proportions across batches. Namely, if we want to achieve indistinguishability of batches in the latent space, a cell type underrepresented in one of the systems must be mixed with a cell type present in the second system^28,30^. We show this with an existing single-cell integration model that leverages adversarial learning, called GLUE^30^, for which it was previously shown to be among the best integration models^41^. This behavior was present in all three datasets, especially when increasing batch correction strength. For example, we observed mixing of delta, acinar, and immune cells in the mouse-human pancreatic dataset; astrocytes with Mueller cells in the organoid-tissue retinal dataset; and adipocyte and ASPC cells in cell-nuclei adipose dataset (Figure 2b, **Supplementary Figure S5**, **Supplementary Figure S6**). In contrast, these cells could be easily separated by our baseline cVAE implementation (Figure 2b**, Supplementary Figure S5**). To address this issue, the authors of the GLUE method aimed to improve the adversarial loss objective by down-weighting the contribution of cells from unbalanced populations. However, this requires prior population identity knowledge, which is in GLUE initialized by a preliminary integration round. Our results show that this strategy does not guarantee good integration performance, which may be explained by biases introduced through imperfect cell cluster prior information. Therefore, adversarial learning may be problematic as a general integration strategy as datasets often have unbalanced cell population proportions due to biological differences or differences in technical protocol capture^42,43^.

## Multimodal priors and cycle-consistency for better integration

To tackle the two above-mentioned challenges of existing methods, that is the joint removal of batch and biological variation and mixing of cell populations with unbalanced proportions across systems, we propose to adapt the prior regularization of the cVAE model and add additional batch correction constraints to the general loss. In particular, we selected the VampPrior^36^ and latent cycle-consistency loss^44^, as they are able to better preserve biological variation.

The VampPrior was initially proposed as an alternative to standard Gaussian prior for generating more expressive latent representations due to the use of multiple prior components^36^ (see methods and Figure 1d). It was previously shown that this also applies to single-cell data^37^, where it increases biological preservation, similar to other flexible priors, such as Gaussian mixtures (GM)^45–47^. Therefore, we here tested whether the VampPrior could be used to achieve a better batch correction and biological preservation tradeoff compared to a cVAE with standard Gaussian prior. We observed that while biological preservation was relatively similar in both models, the model with the VampPrior (VAMP model) had much higher batch correction performance (**Supplementary Figure S1**). As the VampPrior has not previously been used for batch effect removal, we investigated the cause of improved batch correction. We explored the relationship between individual prior components and cell metadata. We assigned individual cells into groups based on the prior component with the highest support for each cell and assessed whether this led to the grouping of cells by either cell types or systems. Prior component groups corresponded more strongly to cell type identity than systems (**Supplementary Figure S7**). This suggests that in contrast to the standard Gaussian prior that attracts cells from different systems and different cell types, using multiple priors allows individual prior components to attract cells from only a subset of cell types that nevertheless originate from both systems, which could explain the improved batch correction at a similar biological preservation score.

To test if batch correction is improved due to the multimodal nature of the VampPrior or its other properties, we compared to a VampPrior model with only one prior component that resembles a normal cVAE, but in contrast to cVAE the prior parameters are learned, and a GM prior model (GMM) as a simpler multimodal prior alternative where prior and posterior distributions are not coupled. We also tested if trainable prior components are key for integration by fixing prior component parameters and inspected if the identity of cells used for pseudoinputs initialization affects the performance.

Across integration data use cases, we find that the batch correction was lower when using only one prior component, compared to two or more for both the VAMP and GMM models (Figure 3, **Supplementary Figure S8**), with the one-component version being more similar to cVAE with standard normal prior (**Supplementary Figure S1**). However, the performance of GMM, but not VAMP, dropped again at very high prior component numbers (N=5000). Therefore, even a relatively small number of prior components (e.g. five) enables good integration while keeping the number of model parameters low. Fixing of prior components had little effect on VAMP performance (Figure 3, **Supplementary Figure S8**, **Supplementary Note S2**) and likewise, the initialization of prior components with cells from a single cell type or system did not affect performance (**Supplementary Figure S9**, **Supplementary Note S2**). This indicates that the number of prior components is key for improved batch correction. Nevertheless, VAMP outperformed GMM both in the achieved batch correction as well as in the robustness to a varying number of prior components, underlining the importance of coupling the prior and the posterior.

**Figure 3:**
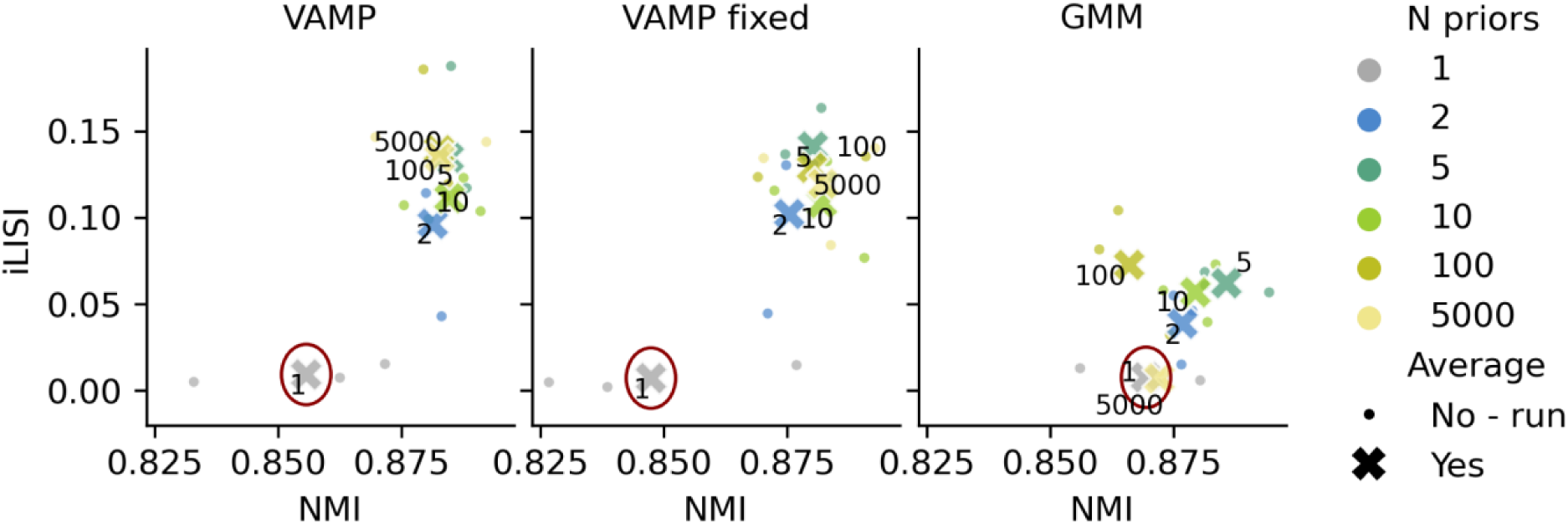
Multimodal priors improve batch correction and biological preservation. Scatter plots of cell-type level biological preservation metric (NMI, higher is better) and batch correction metric (iLISI, higher is better) scores for different models (panels), including a VampPrior model with fixed prior components. Small circles represent individual runs with different seeds and crosses their averages. Points are colored by the number of prior components. The average performances of the models with a single prior component are encircled. Results are shown for the mouse-human data, with other data use cases exhibiting similar trends as presented in **Supplementary Figure S8**.

We propose latent cycle-consistency loss as an alternative to adversarial learning. The cycle-consistency loss directly compares only cells with an identical biological background, that is a cell and its counterfactual representation from another system (Figure 1d). Thus, biologically distinct cell populations are not forced to overlap, unlike in models that aim to align whole system distributions in a cell identity-agnostic manner. While previous publications already used cycle-consistency for integration across modalities (e.g. scRNA-seq and scATAC-seq)^33,35^, they always combined it with adversarial learning, thus overlooking the potential benefits of using cycle-consistency alone.

We validated our hypothesis about cycle-consistency outperforming adversarial learning in the presence of cell populations with unbalanced proportions across systems by implementing a cVAE model with added cycle-consistency loss (CYC model) and comparing it to adversarial model GLUE. For both models, we gradually increased the weight of the loss responsible for batch correction and measured cell type preservation by comparing prior annotation and post-integration clusters with the Jaccard index. When increasing batch correction strength, we observed multiple cell types whose clusters had a lower Jaccard index in the GLUE than the CYC model, thus being merged with other cell types in GLUE (Figure 4, **Supplementary Figure S5**, **Supplementary Figure S6**). Examples are acinar cells mixed with immune cells in mouse-human data, which are not biologically related, adipocytes mixed with adipose stem and progenitor cells (ASPC) in cell-nuclei data, and two glial populations (astrocytes and Mueller cells) in organoid-tissue data. However, some cell types were hard to distinguish for both models, regardless of integration strength. In the mouse-human data, these corresponded to technical doublets, which can be explained by their similarity to individual contributing cell types. Similarly, in the cell-nuclei data neutrophils co-localized with monocytes, likely due to their biological similarity, with both of them being immune cells (**Supplementary Figure S5**). Therefore, none of the models exhibited perfect cell type resolution based on the Jaccard index metric, which is expected as highly similar cell types often require more detailed analysis, including subclustering, for their annotation^48^. Nevertheless, the CYC model was less prone to mix both related and unrelated cell types than GLUE.

**Figure 4:**
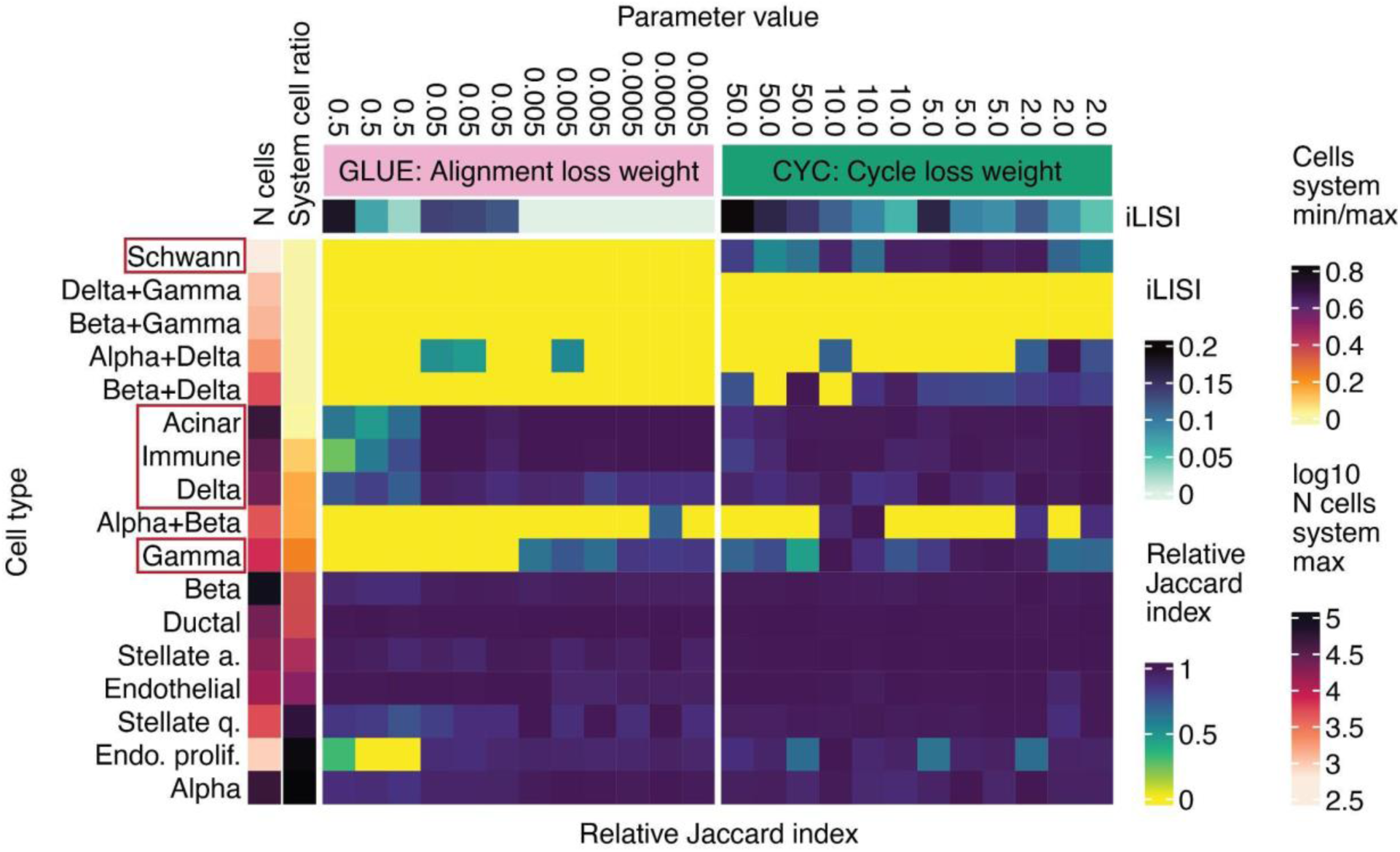
Adversarial learning is more prone to cell type mixing than cycle-consistency loss. Shown is cell type mixing score measured with the Jaccard index between clusters and ground-truth labels (higher is better) and iLISI score. Jaccard index was min-max-scaled per cell type. Results are presented for integration of the mouse-human dataset with GLUE and CYC models with different loss weights of losses that regulate batch correction. Individual cell types are annotated with the number of cells in the more abundant system and the ratio of cells between the less and the more abundant system. Red boxes mark example cell types that are commonly mixed by the adversarial model, especially when increasing batch correction strength. Cell populations representing technical doublets are marked with “+” in their name.

## Integration with the VampPrior and cycle-consistency offers a good tradeoff between batch correction and biological preservation

Our analysis showed that both the VampPrior and latent cycle-consistency loss improved integration performance. Moreover, both approaches are complementary, with VampPrior providing a more expressive latent space than standard Gaussian prior and cycle-consistency loss pushing together batches without incurring cell population mixing in contrast to adversarial learning. Therefore, we propose a new integration approach that combines the two (VAMP+CYC, Figure 5c).

**Figure 5:**
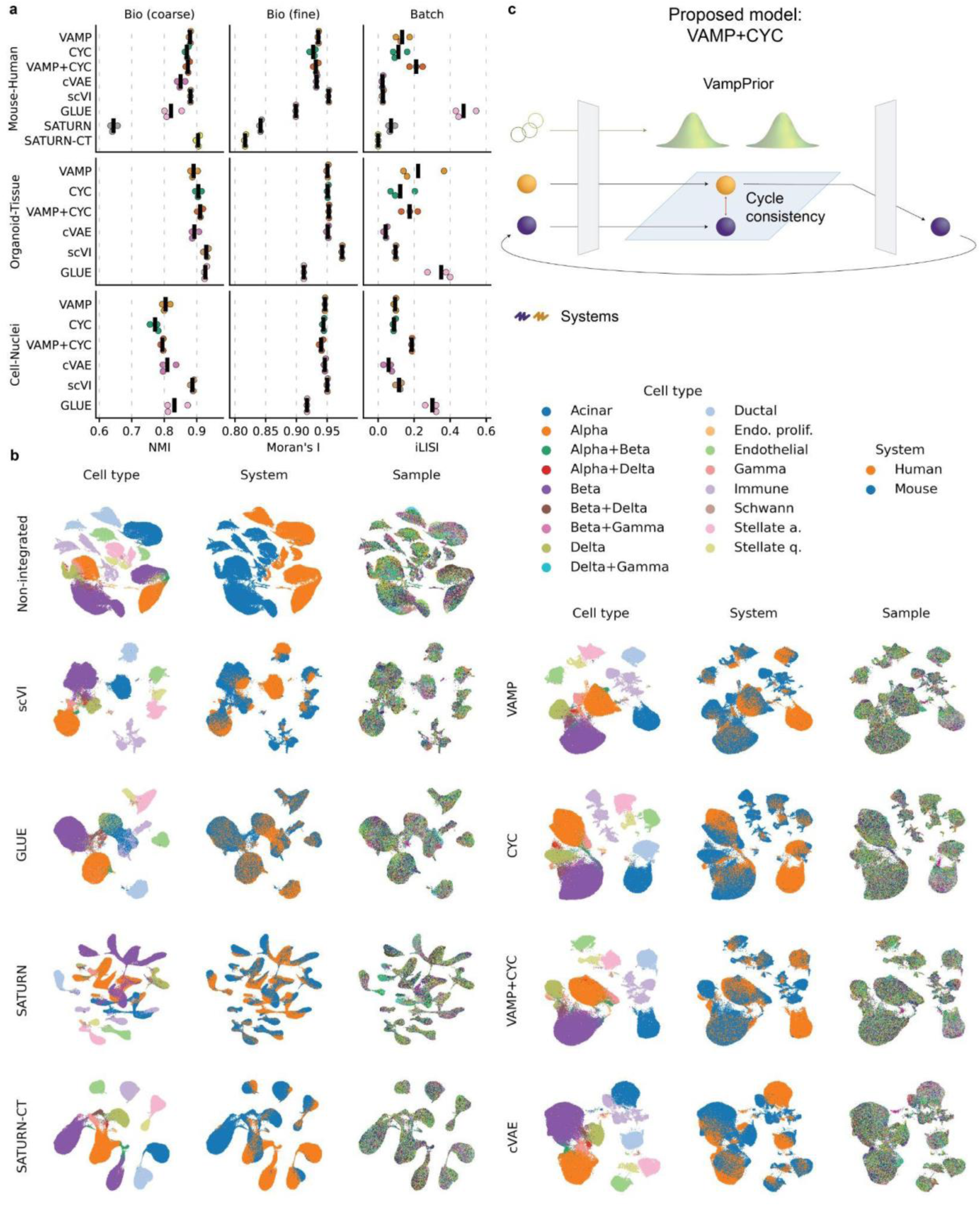
While existing models struggle with over-or under-integration, the VAMP+CYC model achieves a good trade-off between batch correction and biological preservation. (**a**) We show the integration performance of individual models using the best hyperparameter settings, with different integration metrics in columns (larger values indicate better integration performance) and cross-system data cases in rows, showing the average performance (vertical line) of three runs (dots). The results of all hyperparameter settings are in **Supplementary Figure S1**. (**b**) UMAPs of representative runs for the best hyperparameter setting in the mouse-human dataset. A legend for sample colors is not shown due to the large number of samples. UMAPs for the other two data use cases are shown in **Supplementary Figure S5**. (**c**) Schematic representation of the VAMP+CYC model in accordance to the explanation of the VampPrior and latent cycle-consistency loss from Figure 1d.

We compared VAMP+CYC against ablated versions VAMP and CYC, a cVAE baseline, and two established models: scVI^23^, which is a cVAE-based model that models raw expression counts and is among the most popular integration methods, and GLUE, which is an example of a model that uses adversarial learning. We tuned model hyperparameters that directly regulate batch correction, as described in the methods. Every model was run three times with different random seeds to capture random variation in model performance. To complement our integration evaluation via iLISI and NMI, we proposed an additional metric, Moran’s I, aimed to assess fine biological preservation on the sub-cell-type level via preservation of gene expression patterns after integration. Evaluating the fine variation is of special importance as many downstream questions rely on within-cell-type population shifts that arise due to different biological conditions, such as in health and disease.

As hypothesized above, the standard cVAE-based models and the model with adversarial loss could not achieve a good tradeoff between batch correction and biological preservation. The baseline cVAE model had poor batch correction across all tested datasets (Figure 5a), in comparison to other models (adjusted p-value < 0.1 for most comparisons against VAMP+CYC, CYC, VAMP, and GLUE, **Supplementary Table S2**), and the systems were clearly visually separated on the UMAP (Figure 5b, **Supplementary Figure S5**), indicating that it alone is unsuitable for integrating substantial batch effects. In comparison, the cVAE-based model scVI^23^, which is regarded as a state-of-the-art model for scRNA-seq integration^20^, consistently had excellent biological preservation (Figure 5a), in many cases significantly higher than other models (**Supplementary Table S2**). However, it had a relatively low batch correction compared to other models, including VAMP+CYC, CYC, VAMP, and GLUE (Figure 5a), especially in the mouse-human and organoid-tissue datasets that were harder to integrate than the cell-nuclei dataset due to more substantial batch effects (**Supplementary Figure S10**). GLUE had the highest batch correction overall (Figure 5a), which was in most cases significantly higher than other models (**Supplementary Table S2**), and it also had comparable cell type preservation (NMI) to other models in most cases. However, as described above, it led to the mixing of unrelated cell types. It also had lower performance on finer within-cell type variation (Moran’s I) (Figure 5a), which was in all cases significantly lower than other models, except for the below-described SATURN model (**Supplementary Table S2**). As we saw at the cell type level, poor preservation of within-cell-type variation can be explained by the mixing of cell states with unbalanced proportions across systems. Thus, it is likely that the high batch correction of GLUE is not biologically meaningful as it comes at the cost of mixing unrelated cell populations. Integration with adversarial learning is thus less suited for downstream cell state analyses.

In contrast, the VAMP+CYC model, as well as CYC and VAMP models, were able to achieve a better tradeoff between the three integration metrics (Figure 5a) and had across data use cases similar performance characteristics in relation to hyperparameter settings (**Supplementary Figure S1**). They consistently outperformed the base cVAE in batch correction (adjusted p-values < 0.1 in most comparisons, **Supplementary Table S2**) while retaining similar biological preservation. The VAMP+CYC model was able to achieve higher batch correction at comparable biological preservation than the individual CYC and VAMP models (**Supplementary Figure S1**). Namely, the selected VAMP+CYC model had significantly higher batch correction than the two ablated models in the mouse-human and cell-nuclei datasets (**Supplementary Table S2**). In comparison to GLUE, the VAMP+CYC model showed lower batch correction but had higher fine biological preservation (Figure 5a) and was not prone to mixing of cell types with different proportions across systems, as described above. Thus, the VAMP+CYC model is better balanced than GLUE as it has relatively good performance in both biological preservation and batch correction. Overall, these results recommend the VAMP+CYC model as the method of choice.

## Models specific for cross-species integration do not outperform VAMP+CYC

As one of our use cases was cross-species integration, which is complicated by different gene sets across species, we further assessed models that accept different input genes across systems, namely GLUE and SATURN^49^, an example of a model designed specifically for cross-species integration. We ran both models with one-to-one orthologues (OTO) as well as with a flexible orthology (FO) gene set that included different types of orthologues and non-orthologous genes. Integration with FO input genes did not outperform the OTO approach in either GLUE or SATURN (**Supplementary Figure S1**). Furthermore, despite SATURN using protein embeddings to improve gene linking across species, it had significantly lower batch correction than either VAMP+CYC or GLUE (Figure 5a, adjusted p-values < 0.05, **Supplementary Table S2**). Overall, this indicates that a more advanced gene mapping may not be needed for mouse-human integration, as VAMP+CYC was able to outperform even dedicated cross-species integration methods. However, for more divergent species or species with lower-quality genome annotation and orthology information, using FO genes could still be beneficial^19^.

As SATURN requires prior cell type labels for every system, we ran it with clustering-based labels to make it comparable to other models that are likewise not given prior cell type information, which is in practice often not available. Besides that, we also added a version with ground-truth cell type labels (SATURN-CT). However, using ground-truth labels may lead to a positive bias in the NMI metric. Both SATURN and SATURN-CT performed poorly in biological preservation, with SATURN-CT having a bias towards higher NMI scores while retaining low Moran’s I (Figure 5a). The poor preservation of within-cell-type structure was also evident from UMAPs. For example, the numerous immune subclusters observed on the VAMP+CYC embedding were not present after integration with SATURN and SATURN-CT (Figure 5b). The poor biological preservation can be explained by over-reliance on prior cell cluster labels in the contrastive learning objective, which pushes all cells with the same label together, thus neglecting within-cell-type variation (**Supplementary Note S3**).

## High-quality integration is key for downstream biological interpretation

To ensure that a model is of good quality, it must not only have relatively high integration metrics, but it must also be able to provide meaningful biological interpretations. Therefore, we used the integrated embeddings of the evaluated methods for multiple downstream tasks, including cross-system comparison of cell types and conditions as well as the discovery of molecular heterogeneity within individual cell types.

Direct comparison of systems on the integrated embedding requires striking a balance between under and over-integration, that is, to align only biologically related, but not biologically distinct cells. To achieve this, the model’s objective must also reflect this balance by jointly optimizing batch correction and biological preservation (**Supplementary Note S4**). Here, we assess how well models achieve this balance by inspecting individual cases where systems should, in fact, not be completely integrated due to biological differences. We saw that all evaluated models correctly separated retinal pigment epithelial cells between organoid and tissue data^7^ (**Supplementary Figure S5a**). We assessed the preservation of finer cross-system biological differences on the example of Mueller cells, for which it was reported that *RHCG*-expressing fovea cells from primary tissue are distinct from the tissue periphery and organoid cells^7^. This more subtle biological variation was lost in some models, such as GLUE, leading to the co-localization of cells from all three populations (Figure 6a**, Supplementary Figure S11**). In contrast, VAMP+CYC correctly aligned Mueller cells from tissue periphery and organoid samples while separating cells from tissue fovea samples. This indicates its ability to correctly account for biological differences across systems during integration, making it better suited for comparative studies.

**Figure 6:**
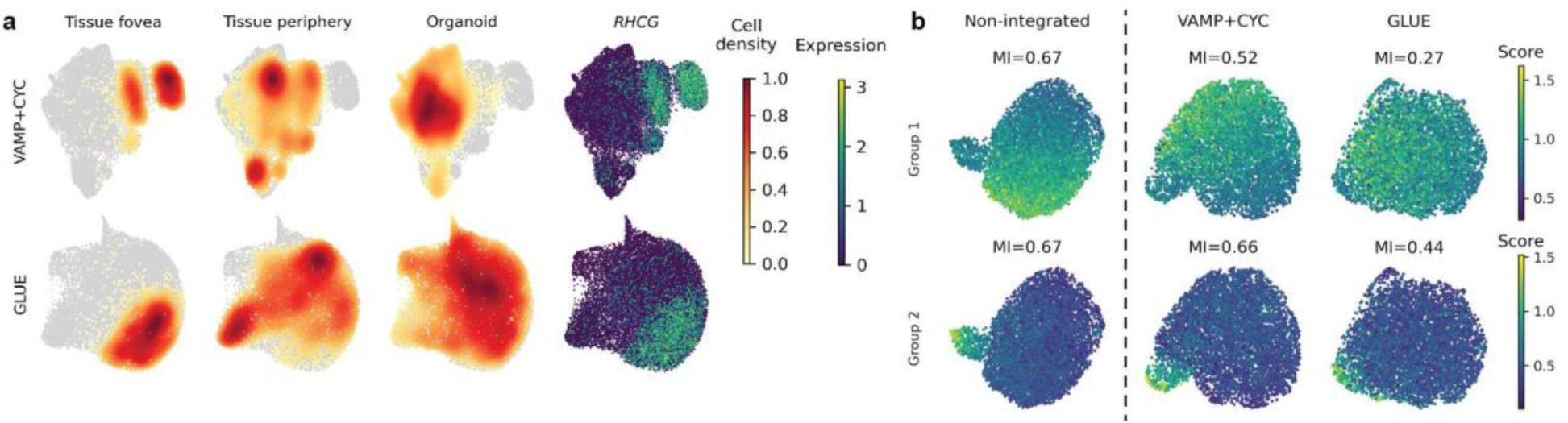
VAMP+CYC empowers post-integration analysis of biological variation. (**a**) UMAPs of Mueller cells based on representative runs for the best hyperparameter setting in the organoid-tissue dataset. The UMAPs are colored by the cell density of organoid and tissue periphery cells, which are expected to overlap, and of tissue fovea cells expressing *RHCG*, which are expected to separate from the other two cell populations. Results for all models are shown in **Supplementary Figure S11**. (**b**) Expression of gene groups known to be heterogeneous within individuals, shown on UMAPs of beta cells from one healthy adult mouse pancreatic sample from the mouse-human data. We compare the heterogeneity before and after integration and quantitatively asses gene group variability with Moran’s I (MI). For every model, one representative run from the best hyperparameter setting was selected (as shown in Figure 5b). Results for all models and remaining gene groups are shown in **Supplementary Figure S12c**.

Another important characteristic of integration is to preserve within-sample variation, which is not related to batch effects. This is key for studying different cell states that arise due to cell specialization or spatial niches. We examined this on pancreatic beta cells, for which within-sample heterogeneity in the expression of multiple gene programs has been previously reported^1^. This heterogeneity was much better preserved by VAMP+CYC than GLUE (Figure 6b, **Supplementary Figure S12c**) and was similar to the heterogeneity observed in the non-integrated data. Therefore, VAMP+CYC has low information loss during integration, empowering post-integration analysis.

## Discussion and outlook

In order to improve the integration of scRNA-seq datasets, in particular in the presence of substantial batch effects such as different protocols, species, or *in vitro* vs *in vivo*, we explored the shortcomings of commonly used cVAE-based approaches for increasing bath correction, namely KL regularization strength and adversarial learning. Both of these models struggled with retaining sufficient biological information when increasing batch correction. To overcome these challenges, we proposed the use of the VampPrior and latent cycle-consistency loss. We performed evaluation in three data scenarios (cross-species, organoid-tissue, and cell-nuclei) and observed consistent model performance characteristics and failure modes, enabling us to make recommendations regarding individual models. We found that our model, which combines the VampPrior and cycle-consistency loss, had an overall good performance in both batch correction and biological preservation as well as enabled truthful interpretation of the integrated embedding, making it our method of choice. To make this model easily accessible to the community, we also implemented it in scvi-tools under the name sysVI.

Based on the findings presented in this work, there are multiple directions of cross-system integration that could be further explored. The VampPrior and cycle-consistency loss could be easily added to other cVAE-based integration tools. Therefore, we urge the method-development community to switch from using batch distribution-matching techniques, such as adversarial learning, to cycle-consistency-based approaches and to replace the standard Gaussian prior with the VampPrior. Combining these approaches with other cVAE extensions may contribute towards achieving the goal of holistic whole-organism and cross-species atlases. The flexibility of the VampPrior holds promise for representation learning on complex datasets with diverse cell types and the cycle-consistency loss further improves the removal of substantial batch effects present in such datasets. To achieve population-wide integration, these two approaches could be combined with scPoli^50^, which enables integration of a large number of batches via learning individual batch embeddings rather than relying on the commonly used one-hot encoding. As a step towards this goal, we included batch embedding in our implementation of sysVI. Furthermore, we here focused on integration strategies that do not require any prior cell type annotation, which is often not available and may lead to biases. Nevertheless, supervised label-aware integration methods such as scANVI^51^, may outperform non-supervised models^20^. Thus, future work could explore the combination of different prior-knowledge-based strategies with the unsupervised techniques proposed here. Additionally, while we heere did not observe a benefit in more complex orthologue mapping for cross-species integration, this may be of greater importance when integrating more evolutionary divergent species. Thus, future work could explore the effect of different gene mapping strategies, both as part of data preprocessing^19^ as well as in the model internally, such as by enabling flexible gene relationships^5,30^ or using gene embeddings^49,52^.

One unexpected finding of this work was that the VampPrior led not only to improved biological preservation, as would be expected, but also to increased batch correction. A similar effect was also observed when using a simpler GM prior. However, the VampPrior showed overall better performance and higher robustness to varying numbers of prior components. Further work will be needed to fully understand the mechanisms of this phenomenon. These results may not only have applications for scRNA-seq integration but also in other domains using cVAE models for covariate effect removal.

While we have shown that VAMP+CYC model enables good cross-system integration in comparison to existing methods, it is not obvious which combination of data analysis decisions will lead to optimal performance and whether integration is the ideal approach at all. For example, here we have focused on evaluating different model architectures but did not analyze alternative data preprocessing decisions that may affect the final result, such as the approach used for selecting features across systems. While previous work assessed some preprocessing options^19,20^, it is unclear if the findings translate to all cross-system data use cases, which can make different assumptions about relationships between systems and their features. Moreover, while our results show that the optimal hyperparameter ranges are relatively similar across data use cases, the preferred setting will depend on the downstream application at hand. For example, annotation transfer across systems would benefit from stronger batch correction, while comparative analysis of systems may require preserving more biological differences in order to assess their true similarity. Furthermore, the coarse biological preservation metric we used (NMI) relies on matching cell type annotation across systems. However, as shown above, sometimes cell types with the same name in different systems are, in fact, biologically distinct and should thus not be aligned, leading to biases in metric interpretation. Instead, one cold use simulated data for evaluation. However, as simulations often cannot fully encompass the complexity of the real data they are likewise not an optimal solution. Therefore, the community would highly benefit from standard benchmarking datasets where proper alignment is more carefully studied, reducing biases in integration evaluation.

While integration eases cross-system comparisons, there are also arguments against performing integration. For example, how strongly to integrate depends on downstream applications and is commonly assessed based on assumptions about the correct system alignment. Therefore, since the final integration is biased toward analysts’ expectations, it may not represent the biological ground truth. The integration also always removes some biological information. This is especially problematic if the integrated systems have substantial biological differences, as much of the biological variation would be lost to enable system alignment. One such example could be the integration of early-stage organoids with adult human tissue, as it is likely that the overlaps between biological functions of cell populations are minimal. In this case, per-system analysis followed by between-system comparison may be preferred. Lastly, recently emerging foundation models claim to obtain batch-free representations of cells via training on a large number of diverse datasets^53,54^, potentially removing the need for future data-use-case-specific integrations. However, as optimal integration strength often depends on the application, it is unlikely that one-size-fits-all models will be able to fully replace data-specific integration. Data-specific tuning of foundation models could improve performance, however this is computationally expensive due to the large number of parameters. Therefore, classical integration models are likely to remain of importance in the foreseeable future.

In conclusion, we proposed an improved strategy for integrating datasets with substantial batch effects that combines a cVAE model with VampPrior and cycle consistency loss. This will ease comparative analyses across a wide spectrum of biological questions, thus better leveraging available scRNA-seq datasets.

## Methods

### Overview of cVAE-based integration approaches

In this study we focused on cVAE-based integration approaches and their extensions that improve integration by modifying the VAE objective (Figure 1d). Here we provide a brief description of these methods.

In cVAE models, the encoder (𝐸_𝜙_) embeds cell expression (𝑥) and batch information (𝑐), into a batch-effect corrected latent representation (𝑧). Then, a decoder (𝐺_𝜃_) reconstructs the expression based on latent representation and batch information. Two opposing losses are used to train the model (combined as 𝐿_𝑐𝑉𝐴𝐸(𝜃,𝜙)_), the expression reconstruction loss that promotes information preservation during encoding and decoding and 𝐾𝐿 loss-based regularization of latent space that encourages information compression towards a Gaussian prior distribution.

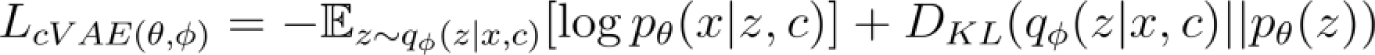

#### Tuning of KL regularization strength in cVAEs

The most straightforward approach for increasing batch correction strength is using a higher 𝐾𝐿 regularization loss weight. This leads to lower preservation of information within the latent space, as it pushes samples’ latent representations towards the Gaussian prior distribution. The information is removed not only for technical or batch variation, which is desired, but as a side effect also for biological variation^26^. To enable good initialization of biological representation the KL regularization loss weight can be gradually increased during training via annealing, as done in scvi-tools^38^.

#### Adversarial loss

Multiple approaches have been proposed for promoting indistinguishability of latent distributions across batches, such as maximum mean discrepancy^27,55^, contrastive mixture of posteriors misalignment penalty^28^, and disentanglement of batch and biology-related latent components^29^. We here chose adversarial learning as an example due to its popularity in the single-cell community^30–32^. The adversarial classification loss (𝐿_𝐴𝐷𝑉_) can be added to the 𝐿_𝑐𝑉𝐴𝐸_ objective to promote indistinguishability of latent embeddings from different batches, with the number of batches 𝐾^30–32^. For this, a discriminator (𝐷_𝑘,𝜓_) is added, which is trained by minimizing classification loss (𝐿_𝐴𝐷𝑉_), while the encoder of the cVAE is trained by maximizing 𝐿_𝐴𝐷𝑉_, opposing the discriminator’s ability to distinguish between batches.

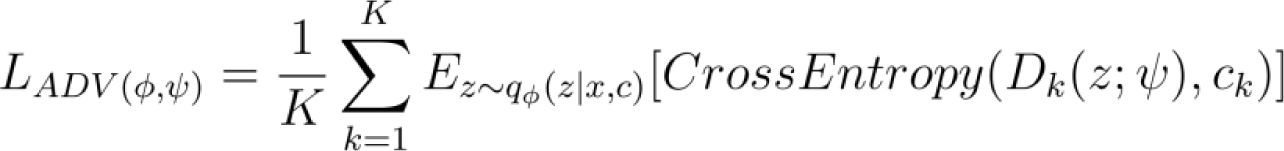

#### Cycle-consistency loss

Latent space cycle-consistency is an alternative to adversarial learning that works by pushing together the latent representation of matched cells from two batches. Cell 𝑥_𝑖_ belonging to the batch 𝑖 is encoded into a latent representation 𝑧_𝑖_. That latent representation is then decoded with batch covariate 𝑗 into the cell 𝑥′_𝑗_, which represents the cell 𝑥_𝑖_as if it originated from the batch 𝑗. The cell 𝑥′_𝑗_is encoded into the latent representation 𝑧′_𝑗_. Finally, the model penalizes the distance between the original representation 𝑧_𝑖_and the cycle-generated 𝑧′_𝑗_via an additional loss component (𝐿_𝐶𝑌𝐶_). Here, different distance minimization losses can be applied, with our choice being the mean squared error (MSE) on standardized data.

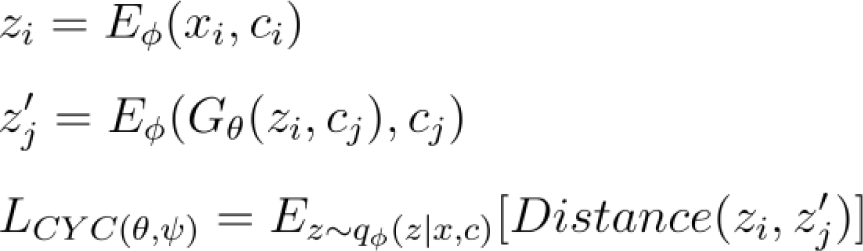

#### The VampPrior

The VampPrior replaces the unimodal Gaussian prior 𝑝_𝜃_(𝑧) with a mixture of trainable Gaussian prior components (with the number of components 𝐿), giving it more representation flexibility. Their prior parameters are not defined in the latent space, as is usually done in cVAEs, but rather in the input cell space as “pseudoinputs” (𝑥𝑝𝑖_𝑙_), which are passed through the encoder to obtain latent representations (mean, variance) used to parametrize the Gaussian components of the prior distribution. This thus directly couples the prior and the posterior. Additionally, the weights (𝑤) of the prior components are likewise learned^36,56^.

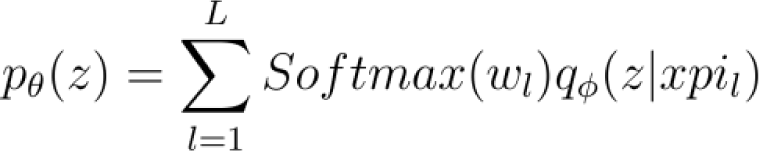

#### Cell type supervision

Another strategy that was previously proposed for improving integration uses prior information about cell type labels for supervised training, thus improving cell type cluster separation. Two main approaches for cell type supervision are contrastive training, where cells from the same cell type are pushed together and cells from different cell types may be pushed apart^49,50^, and classification loss, which ensures that latent space enables good cell type classification performance^51^. Moreover, other approaches were also proposed, such as constraining the cVAE’s prior distribution with prior knowledge about cell types^57^. While in some cases cell type labels can be replaced by unsupervised cell clusters^49^, supervised learning nevertheless depends on the prior information being of high quality. When this is not the case serious integration mistakes may occur, as described above for SATURN. Therefore, supervised approaches are not well suited for new data without annotation. For this reason, we did not focus on cell population supervision in our method comparison.

### Data preprocessing

Data for each of the three use cases was prepared separately as described below. For the mouse-human data, we used pancreatic islet datasets of mouse^1^ (without embryonic and low-quality cells) and human^58^, for the organoid-tissue scenario we used a retinal dataset^7^, and for the cell-nuclei scenario we used adipose dataset^59^ (using the SAT fat type). We obtained published count data and cell annotation for all datasets (see *Data availability* section) and removed unannotated cells. Where necessary we manually curated cell population names to match across studies used within individual integration settings. In each system, we kept genes expressed in more than 20 cells, and in the mouse-human OTO setting we also only kept OTO genes, while in the FO setting, we removed genes without unique gene symbols (required for SATURN). Data was normalized with Scanpy *normalize_total* and log-transformed, and 3000 HVGs (4000 HVGs for organoid-tissue) were selected per system, keeping the union of HVGs across systems, similarly as proposed before^20^. For GLUE we computed a non-integrated principal component analysis (PCA) per system on scaled data using 15 principal components (PCs) and for SATURN we used this data to compute prior clusters per system.

Non-integrated embeddings were computed on the same cells as used for integration evaluation (described below). The normalized expression prepared for integration was standardized per gene, followed by computing 15 PCs, neighbors, and UMAP.

### Evaluation of batch effect strength in unintegrated data

Batch effect strength comparison within and between systems within a data se case was performed by computing Euclidean distances between mean embeddings of cell type and sample groups in the PC space (15 PCs on scaled OTO data). We used only groups with at least 50 cells and cell types where both systems had at least three remaining samples. The significance of differences in distance distributions within and between systems was computed per cell type with the one-sided Mann–Whitney U test.

Batch effect strength analysis of the three data types was performed using average silhouette width (ASW) with systems as the batch covariates for individual cell types of every data use case, using the non-integrated embeddings. We adapted scIB metrics function^20^ so that ASW scores were not reported as absolute values. We used ASW rather than iLISI metric as iLISI was not discriminative enough for the substantial pre-integration batch effects. As the computation was performed on the PCA space with comparable dimension ranges across data use cases, the dimension range bias described in **Supplementary Note S5** does not affect the comparison.

### Integration

We corrected sample and system-level batch effects during integration by adding them to the model inputs as one-hot encoded vectors. The 𝐿_𝐶𝑌𝐶_ and 𝐿_𝐴𝐷𝑉_were computed only on the system covariate. We ran each model with any given hyperparameter setting, as described below, three times with different random seeds.

Our custom cVAE implementation was based on the scVI framework^38^. Unlike in scVI, we used the Gaussian log-likelihood on normalized log-transformed data for reconstruction loss and did not use KL weight annealing. We used the same number of layers and dimensions as for scVI. To regulate batch correction strength we tuned the KL loss weight.

Additional extensions were added on top of our cVAE model. We implemented the VampPrior as described by the authors of the original publication^36^ and did likewise for the GM prior, but with prior components representing points in the latent space. For the VampPrior we initialized prior components by randomly sampling cells from the data and for GM prior we either used sampled data that we passed through the encoder before training or used random initialization with mean in the range [0,1) and variance of one. As in cVAE, we also tuned KL loss weight for the VAMP model. The cycle-consistency loss was computed between the latent representation of a cell (𝑧_𝑐=𝑖_) and its cycle-pair coming from the other system (𝑧′_𝑐=𝑗_) using MSE on data standardized within a minibatch separately for cells and their cycle pairs. In both CYC and VAMP+CYC models we tuned the cycle-consistency loss weight.

All previously published models (scVI, GLUE, and SATURN) were run with default parameters, except for the following changes. For scVI we used *n_layers*=2, *n_hidden*=256, *n_latent*=15, *n_steps_kl_warmup*=1600, and *gene_likelihood*=nb and we tuned *max_kl_weight* (KL loss weight). In GLUE we tuned *rel_gene_weight* (gene graph weights), *lam_graph* (graph loss weight), and *lam_align* (alignment loss weight). In SATURN we used the provided ESM2 protein embeddings and tuned *pe_sim_penalty* (protein similarity loss weight). The number of epochs was set to a fixed value per method and dataset, depending on the number of cells.

### Integration evaluation

We performed evaluations on at most 100,000 randomly selected cells per dataset to reduce the computational cost, except for Moran’s I where cells were selected as described below. We computed neighbors on the latent embedding directly except where we specified that the embedding dimensions were standardized prior to neighbors computation. This data was also used for UMAPs. The non-integrated embedding was computed with 15 PCs on scaled OTO data, using the same set of cells as for the integrated data.

We describe the rationale for metric selection in **Supplementary Note S5**. For LISI and ASW metrics we used implementations from scib-metrics Python package (https://github.com/YosefLab/scib-metrics) and for NMI we adapted their implementation to set a random seed. We computed the NMI-fixed and Jaccard index by first computing Leiden clusters at high resolution (r=2) and then assigning a cell type label to each cluster based on the most common ground-truth label. This annotation was used for comparison to ground-truth labels. For Moran’s I we compared Moran’s I values between non-integrated and integrated data as follows. For the pre-integration Moran’s I computation we kept sample and cell type groups with at least 500 cells and excluded doublets. For each group, we removed genes expressed in less than 10% of the group’s cells, computed 15 PCs and neighbors per group, and used them to compute Moran’s I on all genes. We set a cutoff on Moran’s I per dataset to keep from around a dozen to around 150 genes per group for integration evaluation. On integrated data, we then repeated Moran’s I calculation on the same cell groups and the selected genes. The final score (𝑚𝑖) was defined as a ratio of post-(𝑚𝑖_𝑝𝑜𝑠𝑡_) and pre-integration (𝑚𝑖_𝑝𝑟𝑒_) values averaged across genes (𝐺), samples (𝑆), and cell types (𝐶𝑇).

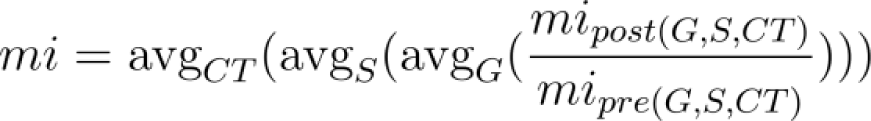

We selected the best hyperparameter setting per model over all tested hyperparameters. For every model, we scaled individual metrics to [0,1] across all runs computed for hyperparameter optimization and then computed the biological preservation score as the average of Moran’s I and NMI and batch correction as iLISI alone. The overall score was computed as a weighted average of biological preservation (weight=0.6) and batch correction (weight=0.4), similar to a previous benchmark^20^. The best hyperparameter setting was selected based on the average overall score across runs and for every hyperparameter setting we selected a representative run for UMAP plots as the run with the median overall score. We observed that optimal hyperparameter ranges were similar across datasets (**Supplementary Figure S1**) and we further discuss considerations for hyperparameter tuning in **Supplementary Note S6**.

We compared the integration performance of all the benchmarked integration methods with their top-performing hyperparameter settings as described above. We used Welch’s t-test followed by multiple test correction per dataset and integration metric with the two-stage Benjamini and Hochberg method. However, a limitation of our analysis is that we always had only three samples per group due to the resource intensiveness of integration benchmarking, reducing the statistical power.

Preservation of gene groups known to be variable within pancreatic islet beta cells of healthy adult mice was evaluated with Moran’s I on the control sample from the mSTZ dataset^1^. We used Scanpy *score_genes* to obtain a single score for every gene group and computed Moran’s I on these scores for every embedding.

### Availability of data and code

The datasets were retrieved from public repositories: GEO (GSE211799), www.isletgenomics.org, Single Cell Portal (SCP1376), and https://cellxgene.cziscience.com/collections/2f4c738f-e2f3-4553-9db2-0582a38ea4dc.

The model and analysis code as well as the conda environments are available at: https://github.com/theislab/cross_system_integration. We implemented our method in scvi-tools package as an external model named sysVI (https://github.com/Hrovatin/scvi-tools/tree/main/scvi/external/sysvi, pull request under review) and provided a tutorial at: https://github.com/Hrovatin/scvi-tutorials/blob/main/scrna/SysVI.ipynb.

## Acknowledgments

We thank members of Theis lab for providing feedback, especially Sabrina Richter. This work was supported by funds from the Helmholtz Association and Helmholtz Munich. F.J.T. acknowledges support from the European Union (ERC, DeepCell—101054957, BetaRegeneration—101054564). Views and opinions expressed are those of the author(s) only and do not necessarily reflect those of the European Union or the European Research Council. Neither the European Union nor the granting authority can be held responsible for them. K.H. acknowledges financial support from Joachim Herz Stiftung via Add-on Fellowships for Interdisciplinary Life Science and support from Helmholtz Association under the joint research school ‘Munich School for Data Science’. M.K. acknowledges support from grants AG081017, NS129032, NS115064, AG074003, NS127187, and AG067151. B.L. acknowledges support from Alana Down Syndrome Center. A.T.L. acknowledges support from ERC Grant DeepCell.

## Author contributions

Conception: K.H. and L.Z.; data curation and illustrations: K.H.; methodology, formal analysis, figures, and original paper draft: K.H. and A.A.M.; supervision: A.T.L., L.Z., B.L., M.K., F.J.T.; funding acquisition: M.K., F.J.T; paper review: all authors.

## Supplementary material

### Supplementary notes

**Supplementary Note S1: KL regularization strength tuning leads to dimension shrinkage, but not batch correction**

Increasing KL regularization strength led, as expected, to stronger batch correction and lower biological preservation. However, we observed that this was not associated with the indistinguishability of cell representations in the latent space, but rather was caused by a reduced number of latent dimensions effectively used in downstream cell graph computation. Namely, stronger KL regularization leads to more latent dimensions having variation near zero (**Supplementary Figure S2**). This occurs as the cell embedding is the predicted mean of the cell’s latent variable representation, as is also done in scVI and GLUE, which is pushed towards zero to match the Gaussian prior when increasing KL regularization. As neighbors graph computation is standardly done with Euclidean distance on non-scaled embedding^60^ this effectively leads to a small number of latent dimensions having a stronger effect on the graph representation. Indeed, after applying standard scaling to the embedding prior to the neighbor graph computation the different KL regularization strengths no longer strongly affected NMI and iLISI (Figure 2a**,b**, **Supplementary Figure S4**, **Supplementary Figure S3**). This indicates that biological and batch information is preserved in the embedding, but is not captured by the standard graph computation protocol. Nevertheless, we observed a drop of within-cell type variation (Moran’s I) on both scaled and unscaled data when increasing KL regularization strength, suggesting that finer variation is nevertheless lost. In contrast, while increasing 𝐿_𝐶𝑌𝐶_loss weight also led to the shrinkage of latent dimensions, the embedding was not sensitive to scaling. We achieved this by computing cycle distances on latent embedding standardized within a minibatch. Thus, while KL regularization strength tuning in standard cVAE merely leads to a lower number of used latent dimensions, thus reducing both biological and batch information at the same time, 𝐿_𝐶𝑌𝐶_actually removes batch effects from the embedding, as indicated by higher batch correction at comparable biological preservation (**Supplementary Figure S1**).

## Supplementary Note S2: Learning of prior components in VAMP

The VampPrior, as proposed in the original publication^36^, contains learnable components and during training the components evolve to be located within data-dense regions (**Supplementary Figure S13**). Interestingly, if prior components are fixed, this does not affect VAMP integration performance, but strongly affects GMM where batch correction drops to a similar level as when using a single prior component (**Supplementary Figure S8**). This is somewhat unexpected, as one would assume that the encoder would be better able to align input representations to fixed prior components when inputs and prior parameters do not lie in the same space, as is the case in the GMM, but not the VAMP.

We also inspected the effect of prior component initialization. VAMP pseudoinputs were initialized with input data. We did not use random initialization as it is challenging to perform appropriate random initialization of pseudoinputs in the input space that would match the input cell distribution and could thus be effectively encoded by the encoder, which is trained to generate input cell representations. In the GMM, the data-based and random prior initialization showed no clear differences (Figure 3). Furthermore, the choice of input cells used to initialize pseudoinputs in VAMP did not seem to play a role, with similar performance when initializing them from a single cell type or system or in a balanced fashion from all cell types or systems (**Supplementary Figure S9**). This indicates robustness against varying prior initialization.

## Supplementary Note S3: Excessive reliance on prior cell cluster labels in SATURN harms cell representation learning

SATURN uses prior cell cluster information to guide cell representation fine-tuning of a pre-trained conditional autoencoder (cAE)-based embedding. This is achieved by maximizing within-species distances between cells with different labels and minimizing cross-species distances between cells that are likely to have the same label. This leads to good cell type level preservation when high-quality cell type labels are available per species (SATURN-CT), however, it does not capture well within cell type information (Figure 5a). This can be explained by the lack of cell expression representation objective in the fine-tuning step, which relies on labels alone. Moreover, the contrastive loss directly pushes cells within cell types together, thus removing within-cluster variation. Thus, it is likely that even an implementation where contrastive loss is used during cVAE training, as in scPoli^50^, may suffer from poor within-cell-type information preservation. In contrast, classification-based loss, for example in scANVI^51^, may be better suited for preserving within-cluster variation as it only enforces that cell clusters separate in latent space, but does not enforce any constraints on within-cluster structure. However, this hypothesis would need to be further tested.

Furthermore, the prior labels must be of good quality to lead to proper cell type separation. While the authors propose that clusters can be used as a source of prior label information, we observed that this led to poor NMI (Figure 5a). Namely, when multiple batches are present within species, as the datasets in our mouse data, this will result in multiple dataset-specific clusters for individual cell types, which will also be reflected in the separation of cells by prior cluster in the final integration (**Supplementary Figure S14**). Similar effects were also observed when prior clusters were too fine or coarse with respect to underlying biological variation. For example, immune cells were separated into three distinct clusters when using cluster-based prior (SATURN), directly corresponding to prior clusters, and were merged into a single cluster when using cell-type-based prior (SATURN-CT). In both cases, the finer cell cluster structure retained within other models was lost (Figure 5b).

## Supplementary Note S4: Excessive reliance on batch covariate information leads to over-integration

Downstream interpretation relies on striking the right balance between aligning cell populations that differ primarily due to batch effects and separating biologically distinct populations. For this, the relevant biological variation and undesired batch effects between systems must be disentangled. This is possible if the batch effects are more consistent across cell types, while system-specific biological variation, such as disease-induced heterogeneity, is cell-type-specific, which may be often the case. Thus, if dissimilar cell types are embedded in the same region of the latent space this will lead to higher reconstruction loss as the batch covariate alone will not be able to explain this variation. However, if reconstruction quality is disregarded after the batch correction step, as described below for scGEN^61^, this may lead to over-integration.

The scGEN model first trains a normal cVAE and then applies an additional batch correction step to every cell type separately. This is done by selecting one batch as the reference and moving all other batches (queries) on top of the reference batch. First, the distance between the reference and query mean latent embedding is computed and then query cell embeddings are transformed by the addition of the distance vector. This will force batch overlap even in the presence of biological differences, making this approach inappropriate for most data use cases as batch and biological effects are usually not orthogonal. For this reason, we did not include scGEN in our benchmark. Nevertheless, we below show that scGEN indeed fails to correctly align biological populations.

All models except scGEN correctly separated retinal pigment epithelial cells from organoid and primary tissue data^7^ (**Supplementary Figure S5a**). In the Mueller cell example, where organoid cells are expected to align with the tissue periphery but not fovea cells^7^, scGEN with samples as batch covariates led to over-integration, placing all samples on top of each other (**Supplementary Figure S11**). The alignment was also not improved by using systems rather than samples as the batch covariate, which instead led to under-integration on the sample level within systems. This also resulted in poor alignment of systems as the alignment with latent space arithmetic was disrupted due to the per-system embedding structure and its mean being strongly affected by sample batch composition rather than being dominated by biology (**Supplementary Figure S5a, Supplementary Figure S11**). This highlights another issue, namely that there is no guarantee that the alignment will be correct on the level of cell subtypes for which corresponding cell annotation is unavailable. This is a consequence of the model never being faced with the task of reconstructing cell expression from the final corrected latent space. For example, we observed some errors in the amacrine subtype alignment, where cells expressing starburst amacrine marker *SLC18A3* did not co-localize in the scGEN embedding while showing clear sub-localization when using the VAMP+CYC model (**Supplementary Figure S15**). Therefore, we deem scGEN inappropriate for integration as its latent space is inherently biologically flawed due to over-reliance on prior information about cell types and batch covariates.

## Supplementary Note S5: Selection of integration metrics

Recently, multiple metrics for the evaluation of integration methods were proposed and implemented in the scIB package^20^. However, we decided to use only a subset that we believe enables a higher-quality evaluation. We observed that metrics that operate on distances directly (such as ASW) rather than on the graph (such as LISI and NMI/ARI) lead to biases when more or less latent dimensions have a high-value range. We observed that when tuning integration hyperparameters, which leads to shrinkage of some latent dimensions (**Supplementary Figure S2**), metrics such as ASW led to inaccurate results that are in the most extreme cases even opposite to graph-based metrics and visually distinct patterns on UMAPs. To validate this, we simulated data (**Supplementary Figure S12a**) where one dimension had a bimodal distribution to represent two distinct groups and 15 additional dimensions represented Gaussian noise unrelated to the two groups. As a starting point, all noise dimensions had the same standard deviation. We then progressively shrank some of the noise dimensions to have a smaller standard deviation, thus corresponding to dimension shrinkage observed when tuning integration hyperparameters. When increasing the number of shrunken noise dimensions, but keeping the group dimension constant, the ASW-based metrics were much more strongly affected than the LISI-based metrics (**Supplementary Figure S12b**). This can be explained by the graph structure being more robust to adding random noise than the underlying distances. As most downstream single-cell analyses are based on graph representations rather than on distances directly, we believe that graph metrics thus more realistically represent data characteristics relevant for analysis. From the group of graph-based batch correction metrics, we decided not to use graph connectivity as it has a poor detection limit^21^ and kBET due to computational costs^21^, and from biological preservation metrics we excluded ARI, as it is computed in a very similar manner as NMI, leading to predominately redundant results (see scIB reproducibility code^20^), and cLISI, as we observed a poor detection limit with the metric being at its maximum in most cases, as also reported before^20^.

While the NMI metric assesses the presence of cell clusters, which are of importance for any downstream single-cell analyses, its interpretation is complicated due to different factors that lead to poor correspondence between embedding clusters and cell type labels. Namely, NMI will be low both when we have merging of multiple cell types, usually caused by too high batch correction, as well as separation of cell types into multiple clusters, which can be caused by different factors. For example, in SATURN individual cell types separate into multiple clusters due to over-reliance on prior labels (**Supplementary Note S3**). In contrast, multiple clusters per cell type may also be a result of poor integration, leading to low NMI even when cell types separate well within batches. This is for example observed in GLUE with small alignment loss weight (0.005 or 0.0005, **Supplementary Figure S1**). Thus, in certain analyses where we wanted to strictly separate between cell type mixing, which is usually associated with excessive batch correction, and the ability to separate cells by cell types, indicating biological preservation within batches, we used an adapted NMI version (NMI-fixed). This metric uses a fixed clustering resolution and then annotates each cluster based on the majority ground-truth cell type. These annotations are used for NMI computation, thus indicating if cell types can be well separated. This corresponds to the ability to do standard scRNA-seq annotation where clustering at high resolution followed by cluster merging is often used to annotate populations that require high resolution to form distinct clusters. However, NMI-fixed cannot detect over-clustering as observed in SATURN, thus we decided to use NMI in most other analyses as it is more generally able to detect if the integrated clusters directly correspond to cell types.

The previously suggested biological preservation metrics^20^ are based mainly on cell type labels, which are commonly available only for cell types, but not for cell states within cell types. Some metrics, such as trajectory, cell cycle, or HVG conservation enable biological preservation evaluation at the sub-cell type level, however, they are often not applicable, due to the lack of a trajectory or cycling cells in the data and integration methods not producing corrected expression values, respectively. Thus, to evaluate how biological variation is preserved within cell types, we adapted Moran’s I biological preservation metric, which was proposed before^1^ to measure spatial covariation of gene expression across an embedding, corresponding to distinct gene expression patterns. We changed the metric to first identify variable genes within individual non-integrated samples and cell type groups and then compare their variation before and after integration for every individual group. These metric modifications enable the selection of more relevant genes that are truly variable prior to integration, remove batch effect biases due to per-sample computation, and enable direct comparison before and after integration. We show examples of Moran’s I values and the corresponding expression patterns for five gene groups known to be variable in mouse healthy adult beta cells^1^ on embeddings produced with different integration models (**Supplementary Figure S12c**). However, as this metric is computed per sample, it cannot detect the preservation of cross-sample patterns or proper alignment across samples. Sample alignment is much more challenging to measure as it usually requires prior knowledge about sample metadata, such as the presence of a developmental trajectory or disease-driven differences across samples within an individual cell type. Therefore, we show sample alignment on a few examples but do not propose a generic metric for it.

## Supplementary Note S6: Effect of hyperparameter tuning on the performance of different models

One important characteristic of integration models is the ability to tune hyperparameters regulating batch correction to achieve sufficient integration for downstream analyses depending on the batch strength. At the same time, increasing batch effect correction often reduces the preservation of biological information^20,55^, necessitating finding a good tradeoff. We observed that some models did not offer enough flexibility in batch correction tuning or contained an overwhelming number of possible parameter combinations to be accounted for. Furthermore, some hyperparameters weren’t monotonically associated with batch correction, in contrast to our expectations.

In the scVI and VAMP models, stronger KL regularization resulted in lower biological preservation. However, iLISI did not increase consistently with increasing KL regularization strength and remained relatively low in all settings (**Supplementary Figure S1**, **Supplementary Figure S16**). This biological preservation-batch correction tradeoff makes this hyperparameter-model combination less favorable. In contrast, while increased 𝐿_𝐶𝑌𝐶_weight in CYC and VAMP+CYC models was also negatively correlated with biological preservation, this was somewhat less prominent than when tuning KL regularization strength, and the positive correlation with batch correction was also higher (**Supplementary Figure S1**, **Supplementary Figure S16**). Furthermore, 𝐿_𝐶𝑌𝐶_ was also able to achieve higher iLISI than the tuning of KL regularization strength alone. Altogether, this makes 𝐿_𝐶𝑌𝐶_, in comparison to KL regularization strength, a better candidate for custom batch correction tuning of cVAEs, both when using a Gaussian prior and the VampPrior.

Multiple hyperparameters of GLUE (adversarial alignment loss weight, gene graph loss weight, and gene graph edge weight) affected integration performance in different ways (**Supplementary Figure S1**). We observed that increased alignment loss weight, which should increase adversarial batch correction, had an optimal iLISI range (around 0.05 in all datasets) and thus did not increase batch correction beyond a certain point, similarly as described for KL regularization in the scVI and VAMP models. Note that the low NMI at low alignment loss weight is not caused by over-integration but rather by the separation of cells across the system, thus leading to multiple clusters per cell type, which then do not match up with the cell type labels (**Supplementary Note S5**). Furthermore, decreasing gene graph loss weight and gene graph weights led to higher iLISI, with low gene graph weights leading to the highest iLISI overall, at the expense of biological preservation (**Supplementary Figure S1**, **Supplementary Figure S16**). This could be explained by the role of the gene graph in biological supervision, as it connects the gene embeddings and subsequently also cell embeddings from individual encoders. If the gene graph constraints are reduced, the gene correspondence between systems can be re-interpreted to increase batch correction. While this may be beneficial in moderate amounts, for example for cross-species integration, it could potentially, in extreme cases, lead to biologically false correspondence between genes and consequently cells. Furthermore, weaker graph constraints may lead to lower graph loss, with lower graph loss resulting in relatively higher importance of the alignment loss during training. Overall, the multitude of losses and their interactions in GLUE that affect integration performance complicates the selection of the hyperparameter set to tune.

Lastly, as SATURN didn’t offer many loss parameters to be optimized, we tuned only protein similarity loss weight that regulates the reliance on prior-defined protein embedding of genes. Despite a wide range of the tested values (0.01-10.0), we did not observe major variations in biological preservation or batch correction.

It is important to consider that the observed hyperparameter-batch correction patterns hold only in the tested hyperparameter ranges. Nevertheless, the tested hyperparameter ranges were selected so that values outside of them would likely lead to inadequate biological preservation or batch correction, making them unsuitable for integration.

## Supplementary methods

### Simulation for comparison of graph and distance-based metrics

We simulated two groups of points (n=100 per group) from normal distributions with means of zero and one, respectively, and variance of one. We added 15 additional dimensions coming from a standard normal distribution, representing noise dimensions that do not separate between the two groups. ASW and LISI (both biological preservation and batch correction variants) were computed on all 16 dimensions while the size of 0 to 15 noise dimensions was divided by ten to push them towards zero. The simulations were repeated ten times.

### Integration with scGEN

We ran scGEN with default parameters, except for *kl_weight=0.1*. We used either samples or systems as the batch variable since the model does not allow for multiple batch variables.

## Supplementary figures

**Supplementary Figure S1:**
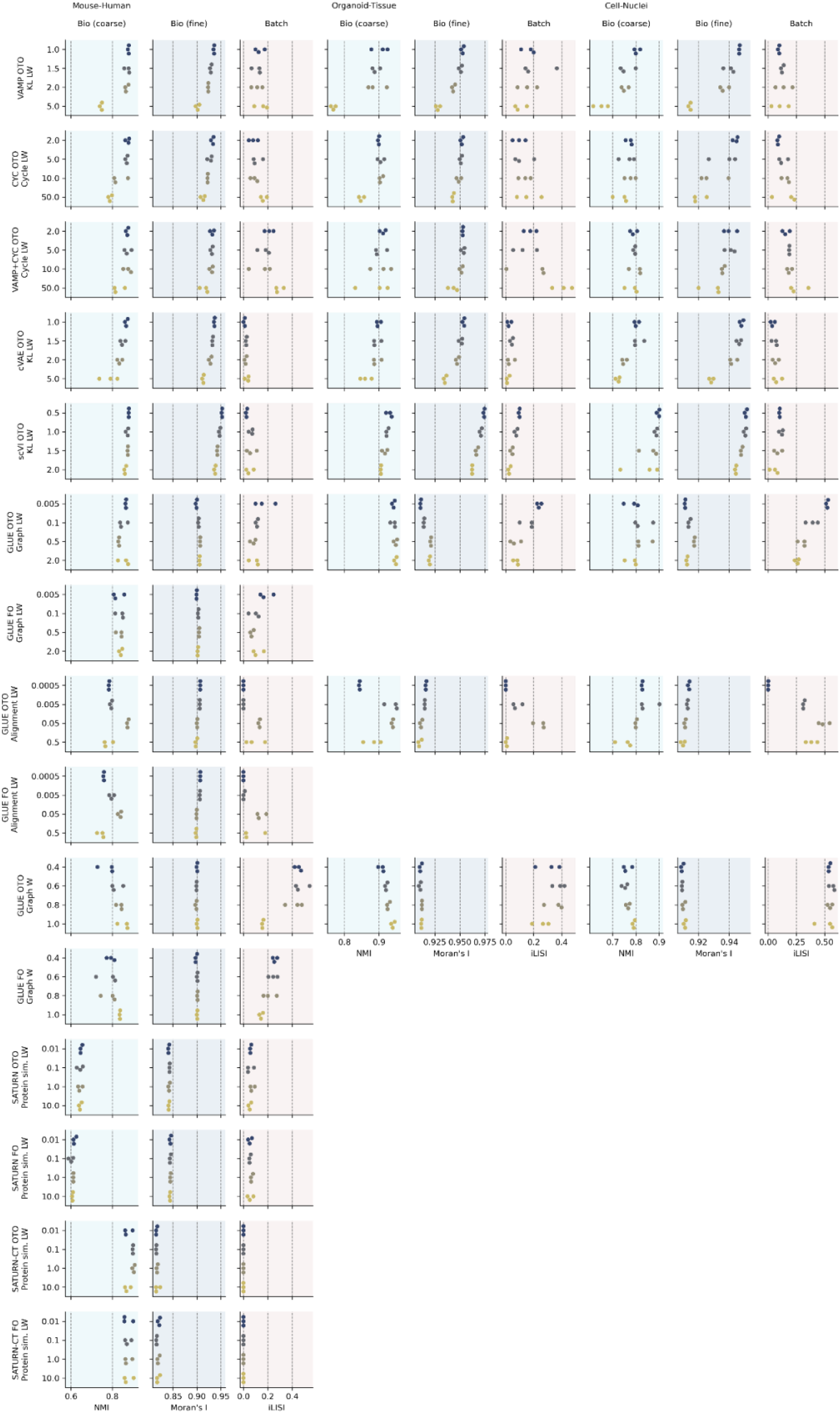
Integration performance across models and different model hyperparameters. Rows show different models and tested hyperparameters (y-axis values), columns show per-dataset integration metrics, and dots represent individual runs with different seeds.

**Supplementary Figure S2:**
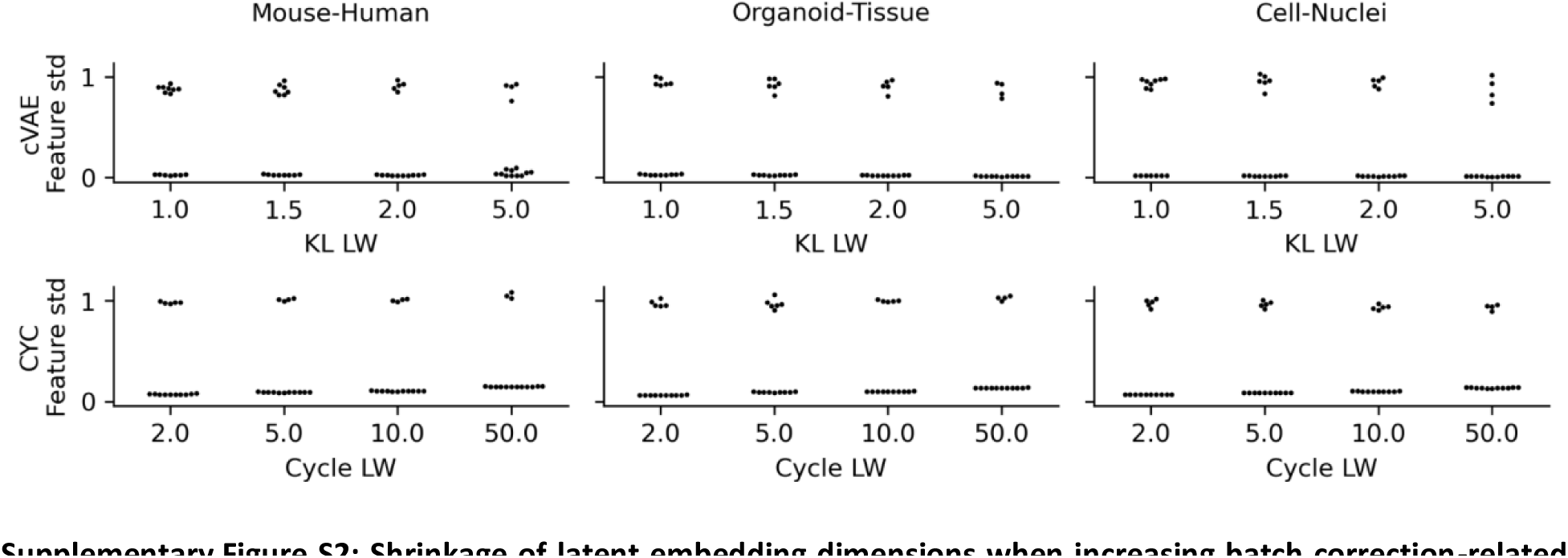
Shrinkage of latent embedding dimensions when increasing batch correction-related hyperparameters. Standard deviations (std) of individual embedding dimensions (dots) obtained with different hyperparameter values (x-axis, shown for runs with seed=1) for cVAE and as a comparison CYC (rows) across datasets (columns).

**Supplementary Figure S3:**
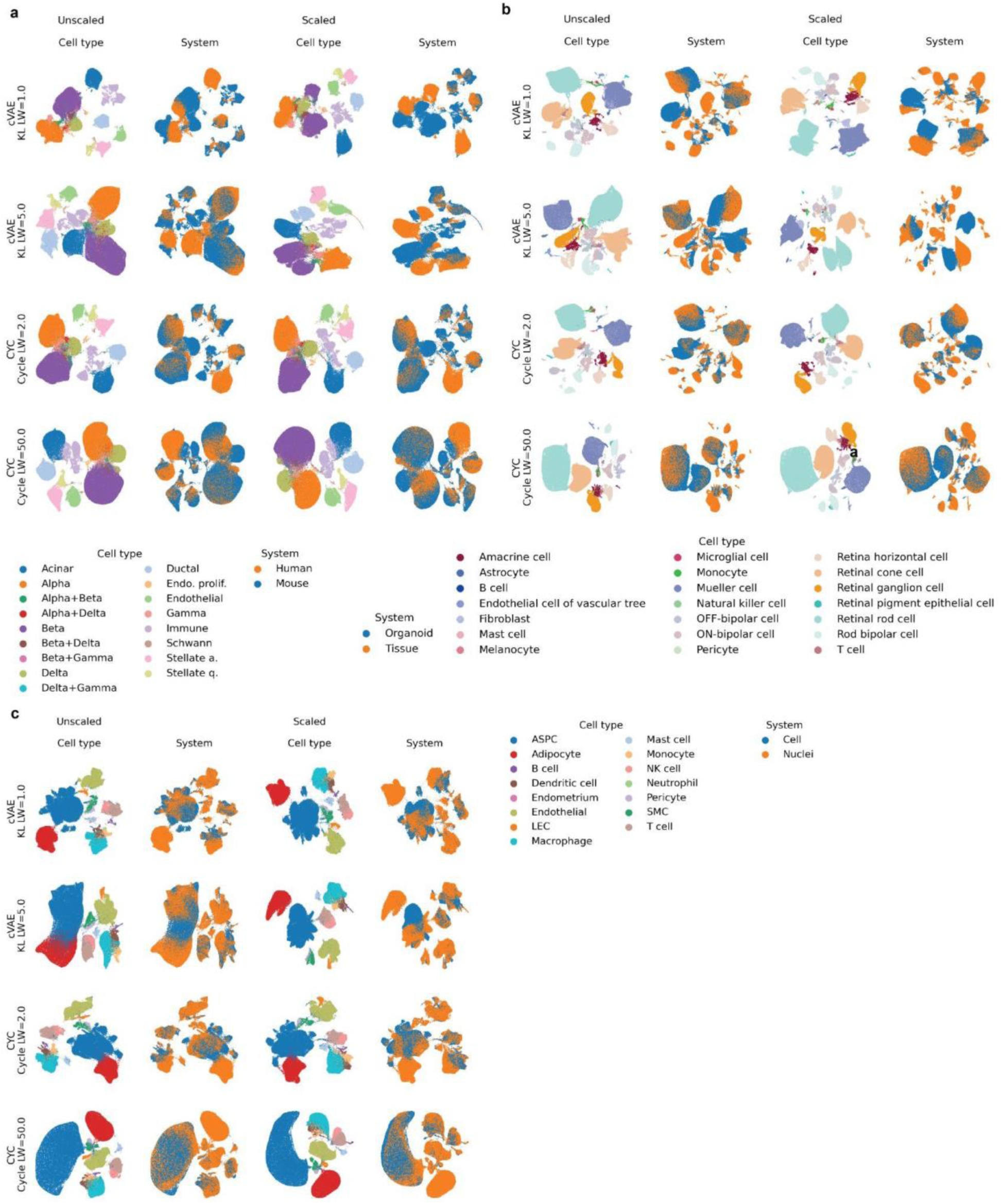
UMAP of scaled and unscaled embeddings. Embeddings produced with cVAE and CYC models with low and high batch correction strength hyperparameter values (shown runs with seed=1). Datasets: (**a**) mouse-human, (**b**) organoid-tissue, (**c**) cell-nuclei.

**Supplementary Figure S4:**
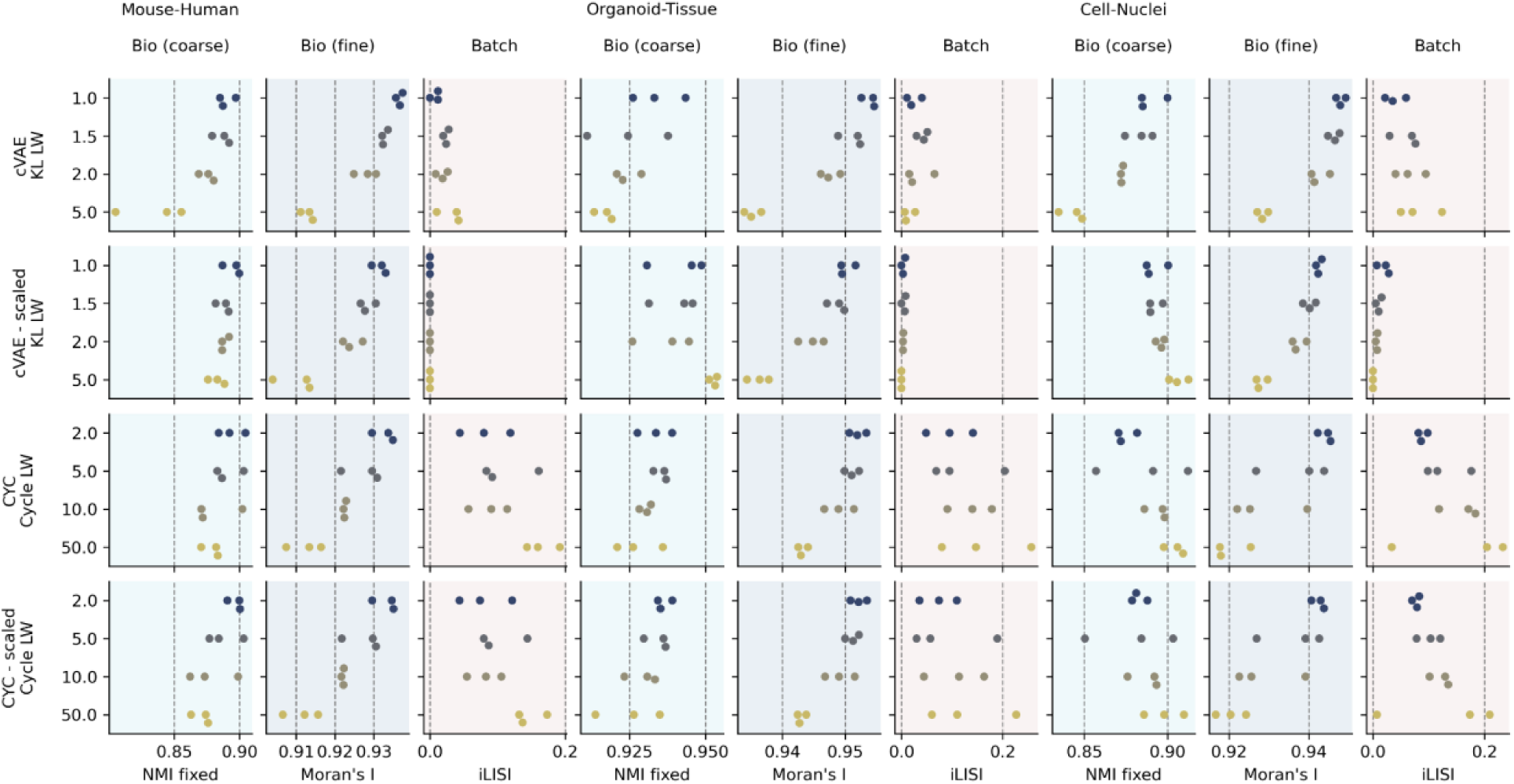
Effect of scaling cVAE integration embedding on integration metrics. Integration metrics of datasets (x-axis) computed using scaled and unscaled embedding from cVAE and for comparison CYC, shown across different values of hyperparameters used for tuning batch correction (y-axis).

**Supplementary Figure S5:**
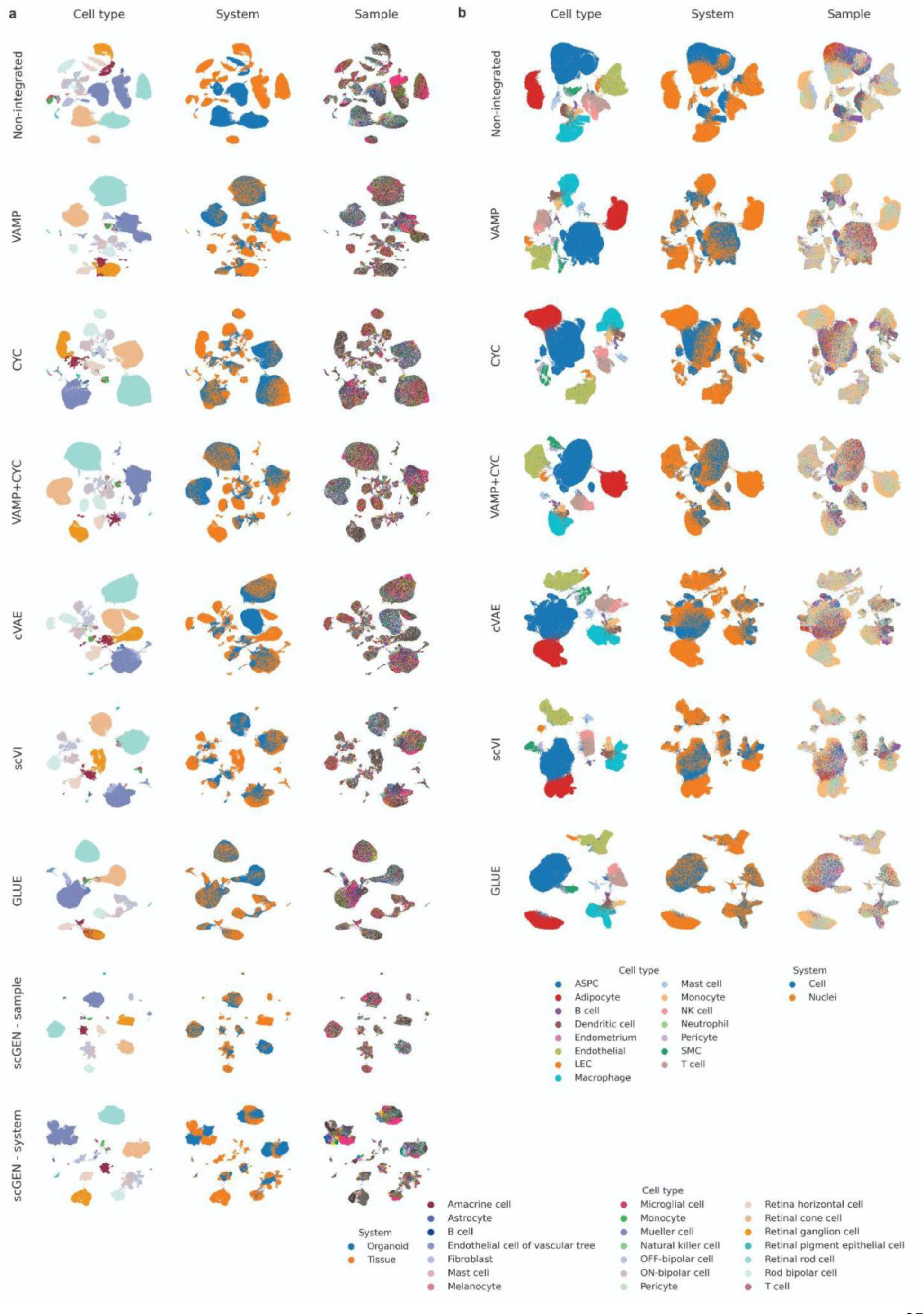
UMAP embeddings of representative runs for the best hyperparameter setting of every model. Shown for datasets (**a**) organoid-tissue and (**b**) cell-nuclei. To (**a**) we also added two example runs for scGEN with samples or systems as batch covariates. A legend for sample colors is not shown due to the large number of samples.

**Supplementary Figure S6:**
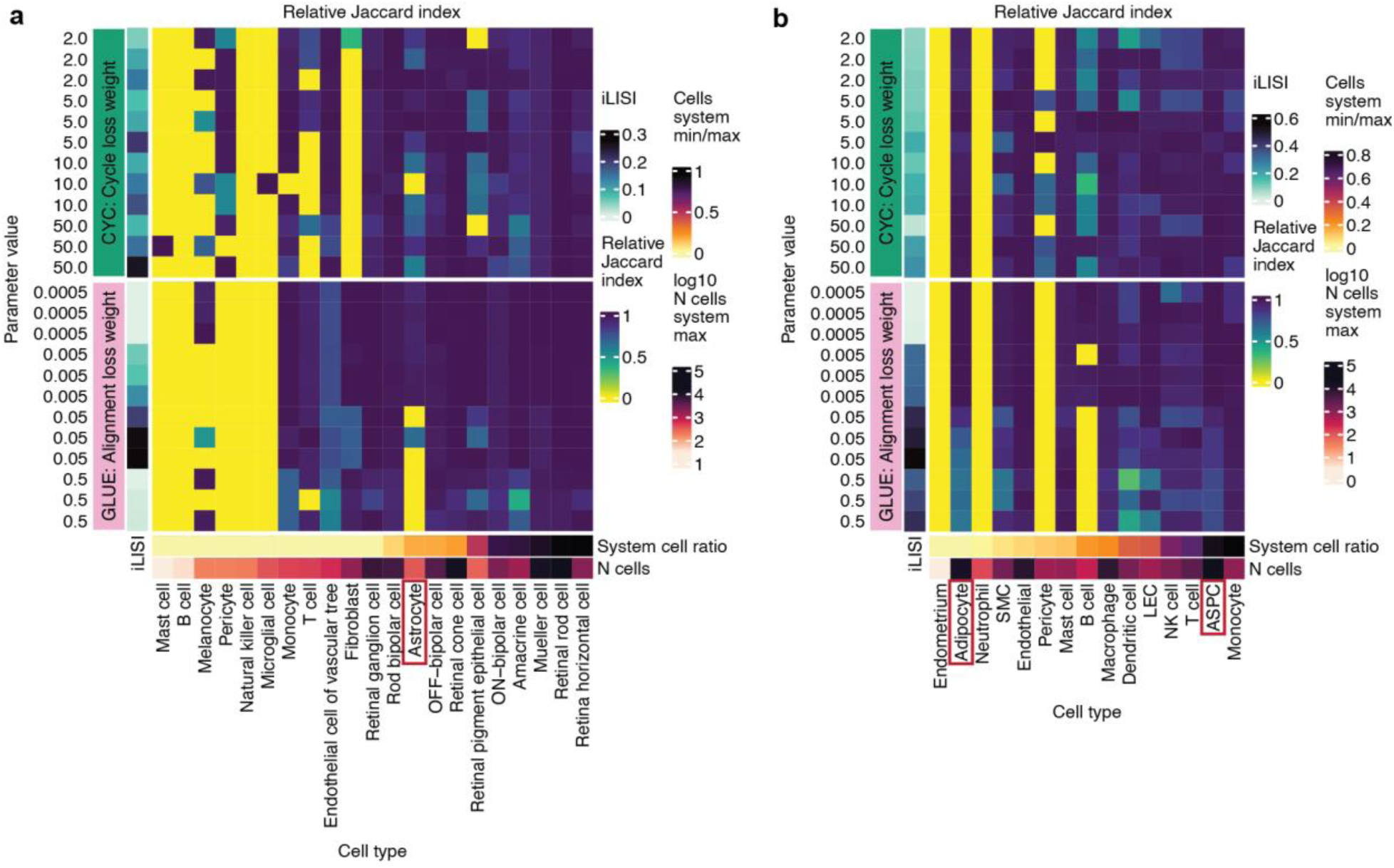
Adversarial learning is more prone to cell type mixing than cycle-consistency loss. Shown is the cell type mixing score measured with the Jaccard index between embedding clusters and ground-truth labels; the score was max-scaled per cell type. Results are presented for integration of the (**a**) organoid-tissue and (**b**) cell-nuclei datasets with GLUE and CYC with different loss weights (LW) of losses that regulate batch correction. Individual cell types are annotated with the number of cells in the more abundant system and the ratio of cells between the less and the more abundant system. Red boxes mark example cell types that are commonly mixed by the adversarial model, especially when increasing batch correction strength.

**Supplementary Figure S7:**
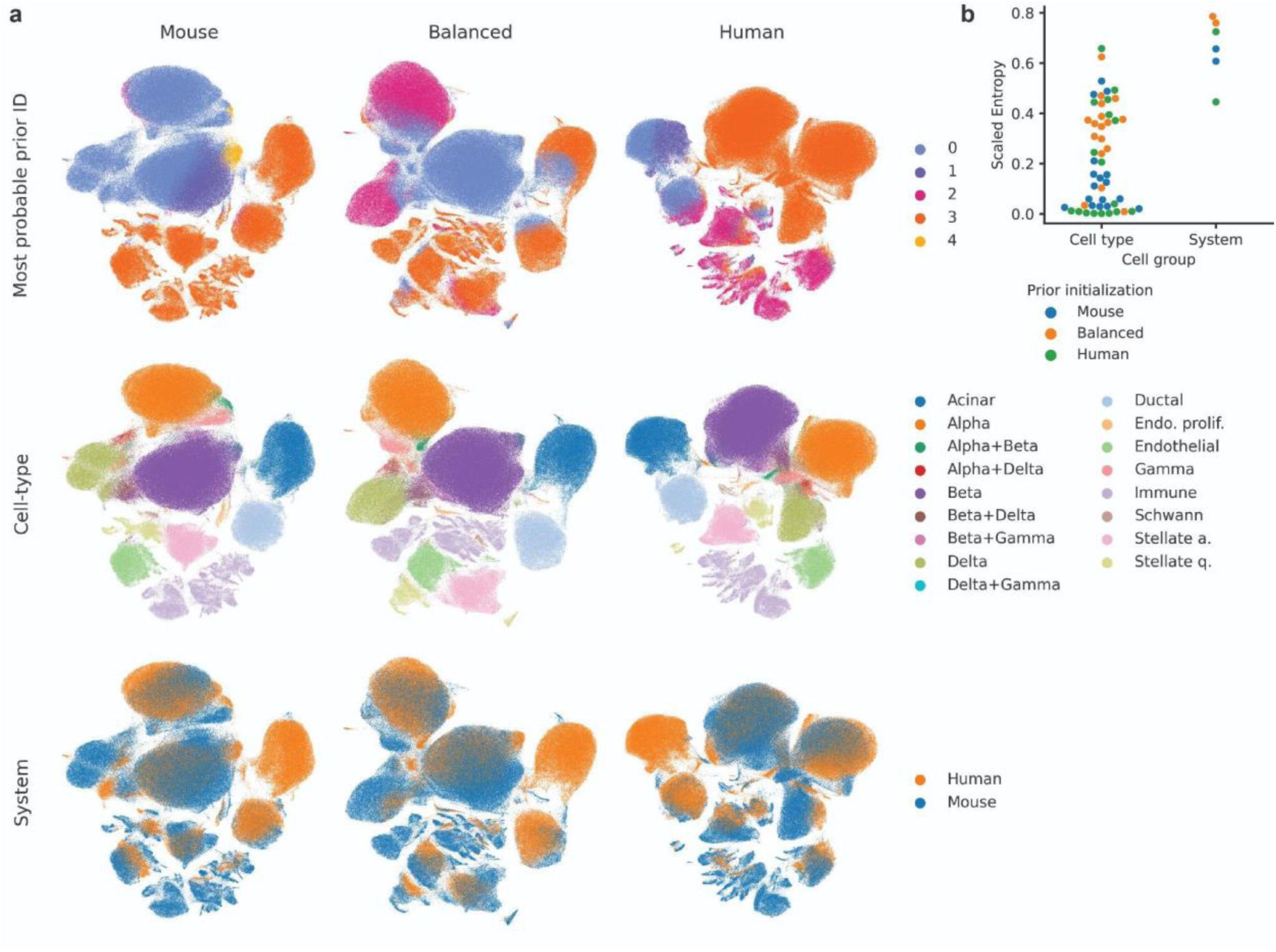
Individual VampPrior prior components co-localize with cell types rather than systems. **(a)** Cells are colored by the prior component (N=5) with the highest probability, cell type, and system (rows) on embeddings generated with different prior initializations (columns). Priors were initialized either by sampling cells from a single system or in a balanced manner from both systems. Shown are UMAPs of the VAMP integration of the mouse-human dataset. (**b**) Entropy of the most probable prior assignment within every cell type or system (dots) corresponding to (**a**), with colors indicating runs with different prior initializations. Entropy was scaled by the maximal possible entropy.

**Supplementary Figure S8:**
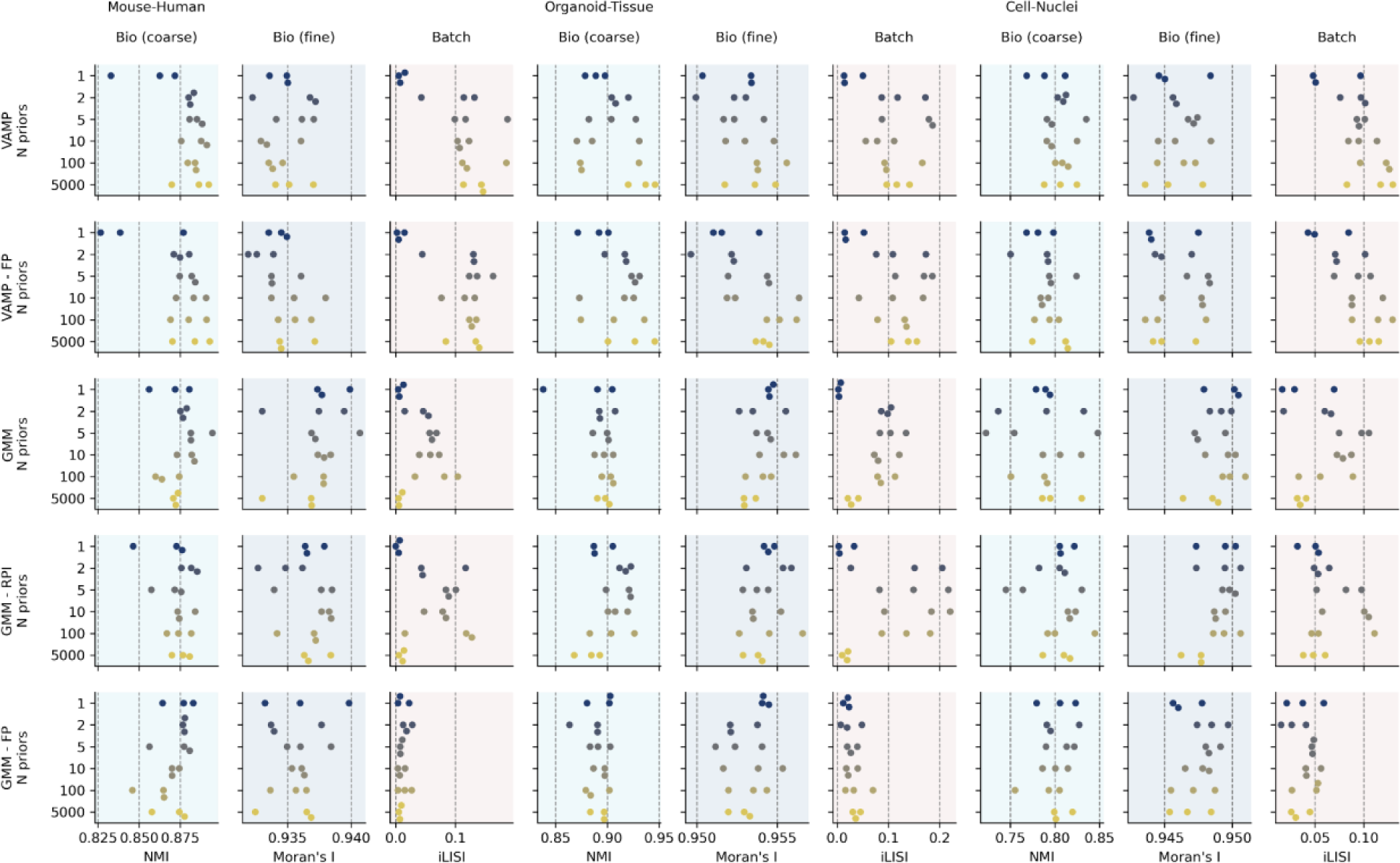
Integration performance of VAMP and GMM with different prior component settings. Rows show different models and prior settings, including the number of prior components (y-axis values), columns show per-dataset integration metrics, and dots represent individual runs with different seeds for each setting. FP - fixed prior, RPI - random prior initialization.

**Supplementary Figure S9:**
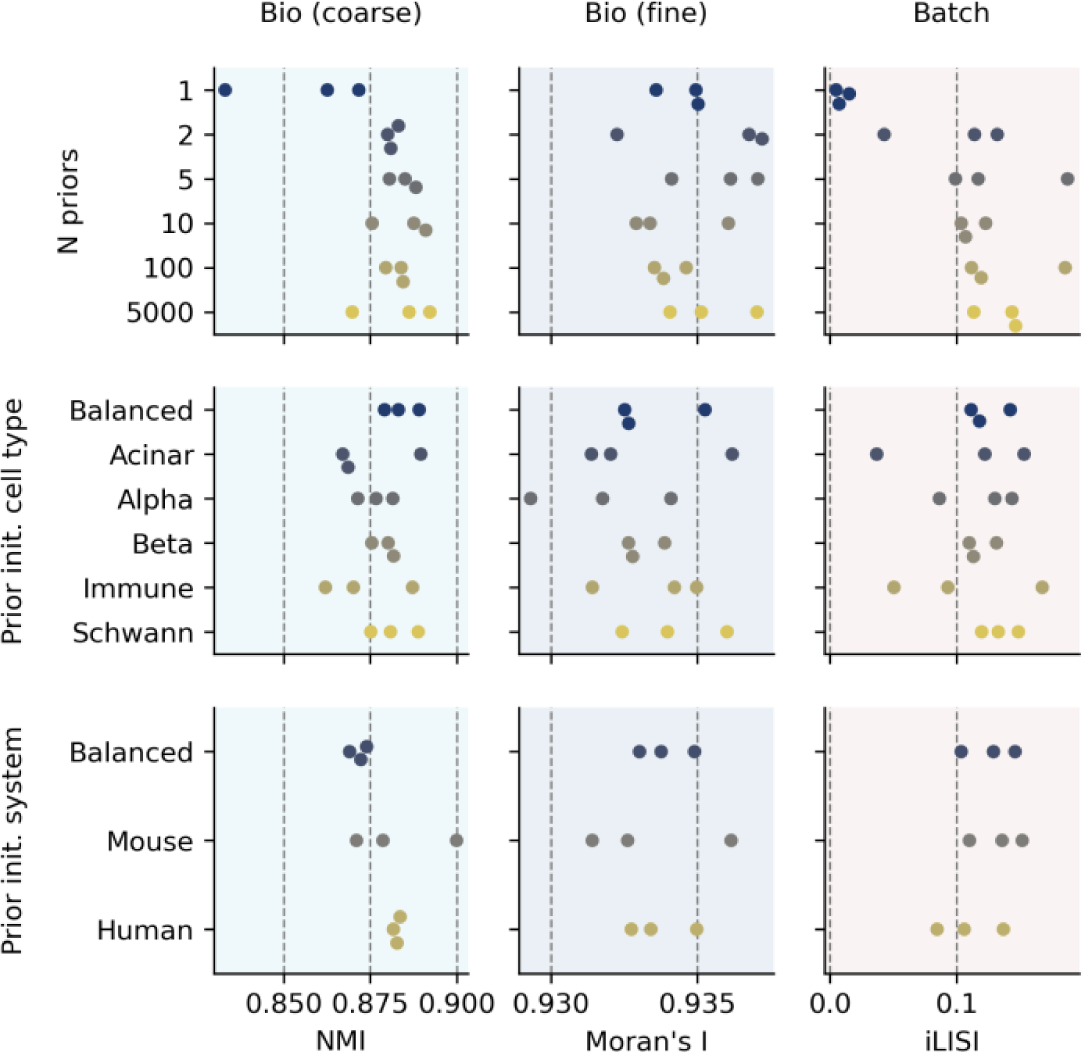
Initializing pseudoinputs from all cell types or both systems or only one cell type or system does not have a major effect on the integration performance. Rows show different prior initializations, with the top row having random initialization with a different number of prior components (y-axis), and the other two rows being initialized with either one prior component sampled from each cell type or system (“Balanced”) or from a single cell type or system with the number of prior components equal to the number of cell types (N=17) or systems (N=2), respectively. Columns show integration metrics for the mouse-human dataset and dots represent individual runs with different seeds.

**Supplementary Figure S10:**
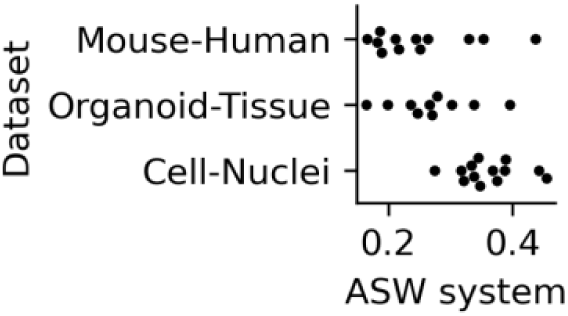
Between-system batch effects strength prior to integration. Shown are ASW batch values computed with systems as batch covariates for individual cell types (points) in every dataset, with higher scores indicating stronger system mixing. The distribution of ASW batch scores is significantly higher in cell-nuclei dataset compared to mouse-human (p-value = 2.1e-03) or organoid-tissue (p-value = 3.9e-03) dataset, with the latter two not differing significantly from each other (p-value = 3.7e-01). This indicates that batch effects in the cell-nuclei dataset are relatively weaker, as can also be seen in the non-integrated UMAP plots (Figure 5b, **Supplementary Figure S5**).

**Supplementary Figure S11:**
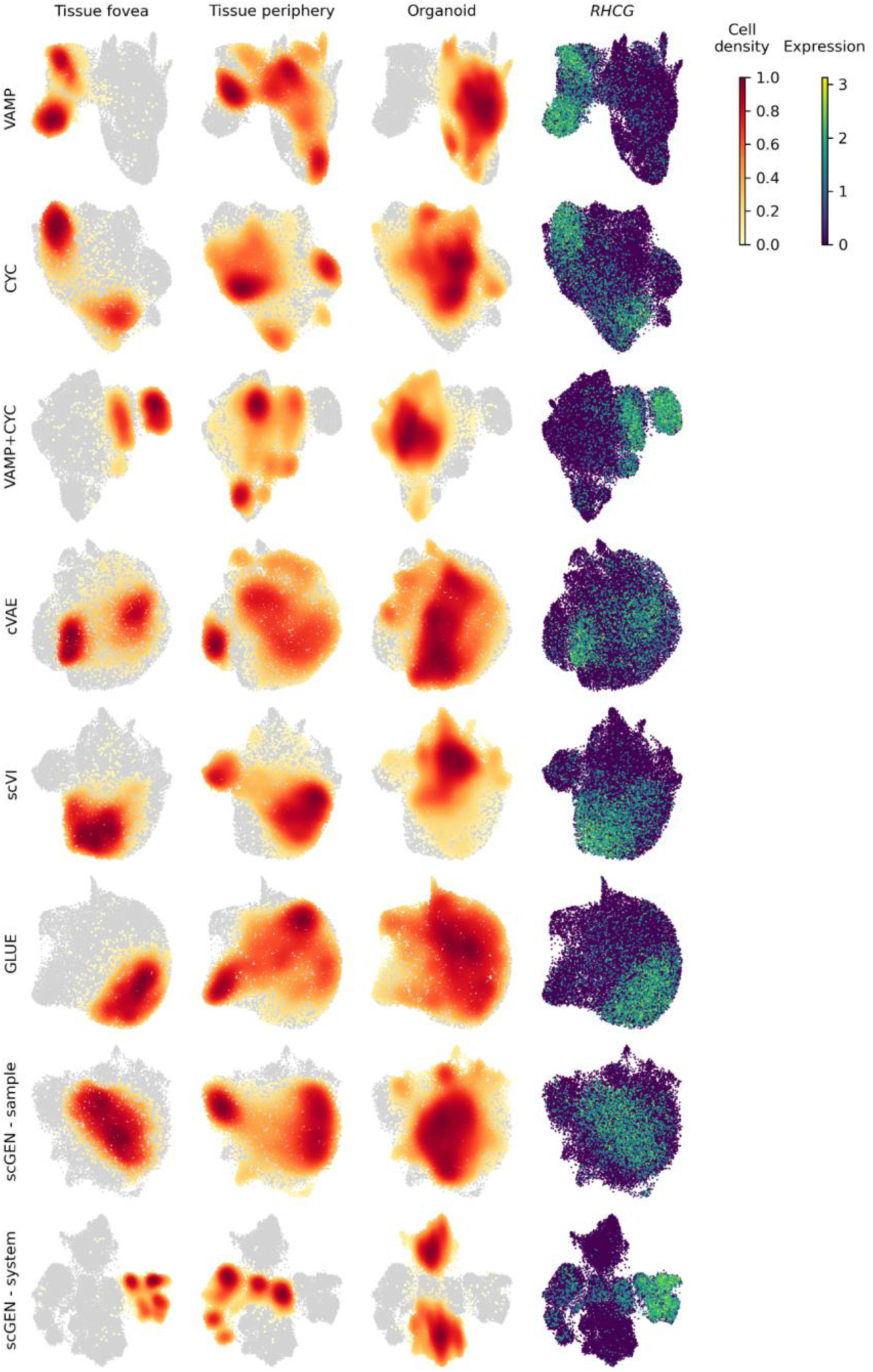
Sample alignment in integrated Mueller cells. Plotted are UMAPs of Mueller cell subset from the organoid-tissue dataset based on representative runs for the best hyperparameter setting of every model. We also added two example runs for scGEN with samples or systems as batch covariates. The UMAPs are colored by cell density in organoid and tissue cell groups and expression of foveal marker *RHCG*.

**Supplementary Figure S12:**
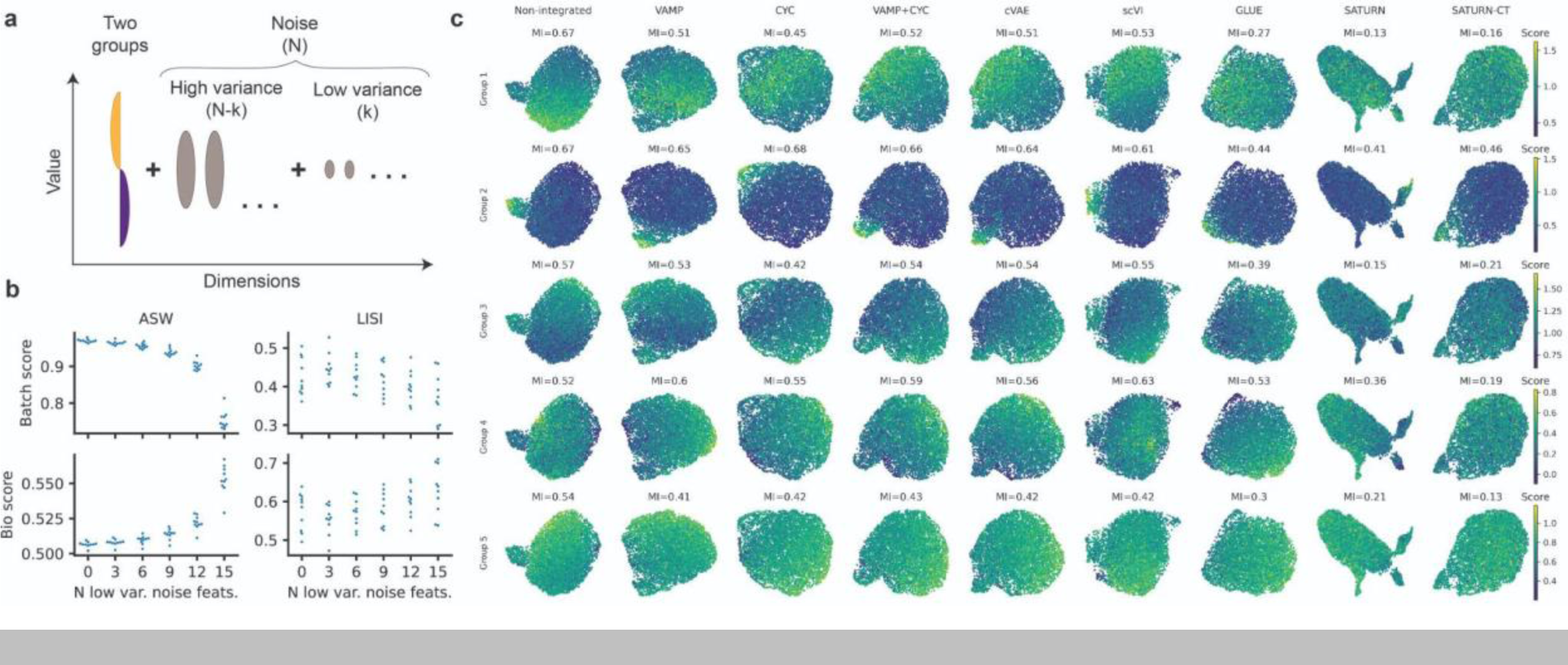
Evaluation of integration metrics. (**a**) Overview of the data simulation process used for comparing distance and graph-based metrics. (**b**) Graph-based metrics are less affected by the value range of the noise dimensions than distance-based metrics. Shown are biological preservation and batch correction scores based on ASW or LISI when 0 to 15 of the noise dimensions are shrunk to have smaller variance. (**c**) Examples of Moran’s I scores and corresponding expression patterns on UMAPs for five gene groups known to be variable in healthy adult mice. For every model, one representative run from the best hyperparameter setting was selected (as shown in Figure 5b) and cells were subsetted to one healthy adult mouse pancreatic sample, used for computing UMAP and Moran’s I.

**Supplementary Figure S13:**
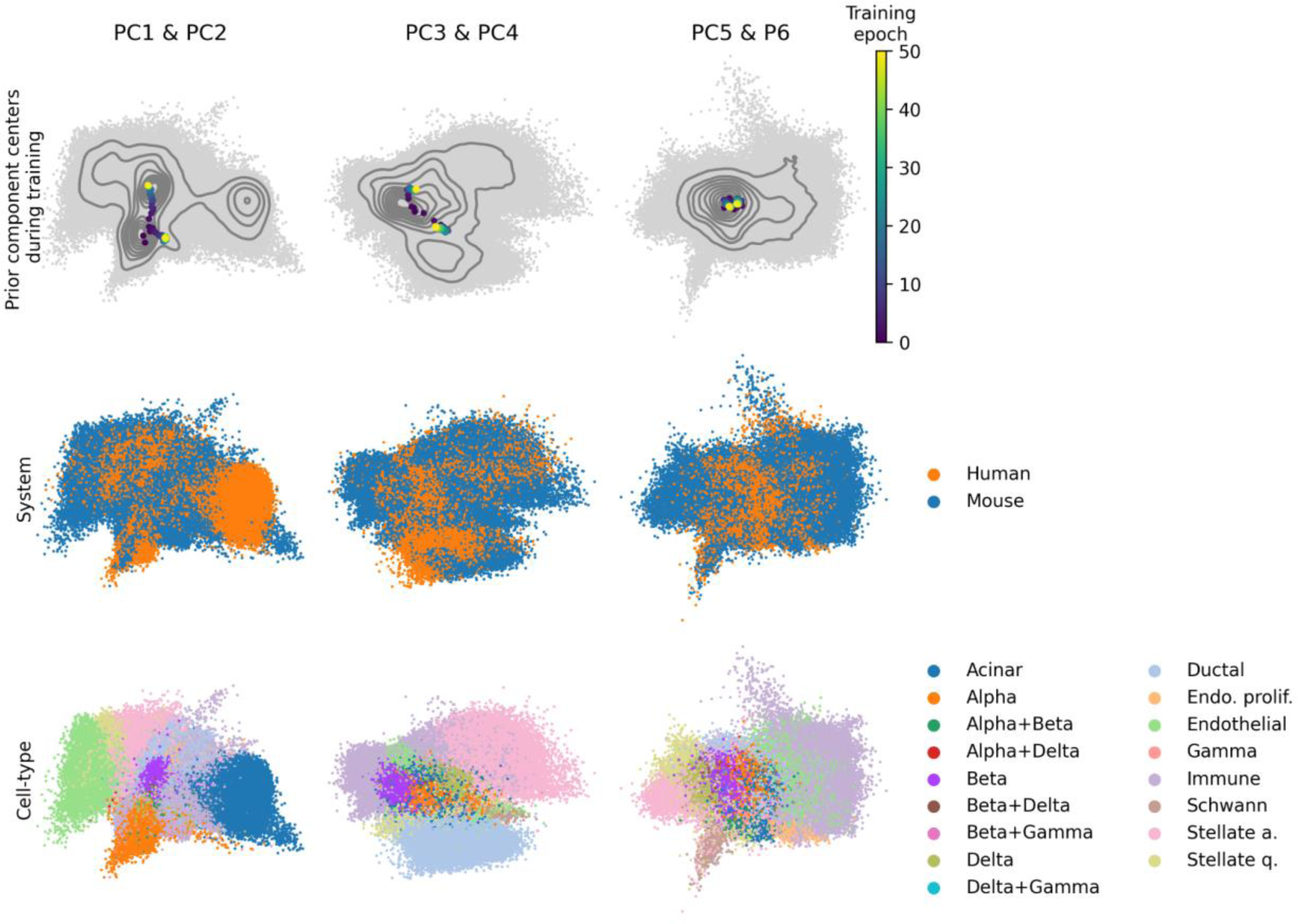
The update of the VampPrior components during the training. The top row shows how the VampPrior components’ (N=2) mean parameters (colored) are updated during the training in the space of the first six PCs (pairwise plots in columns) computed on the model’s latent space. The contour lines represent the density of the input cells (every 10th quantile) and the gray points individual cells. The middle row shows the system covariate and the bottom row shows the cell type. Shown are UMAPs of VAMP integration of the mouse-human dataset.

**Supplementary Figure S14:**
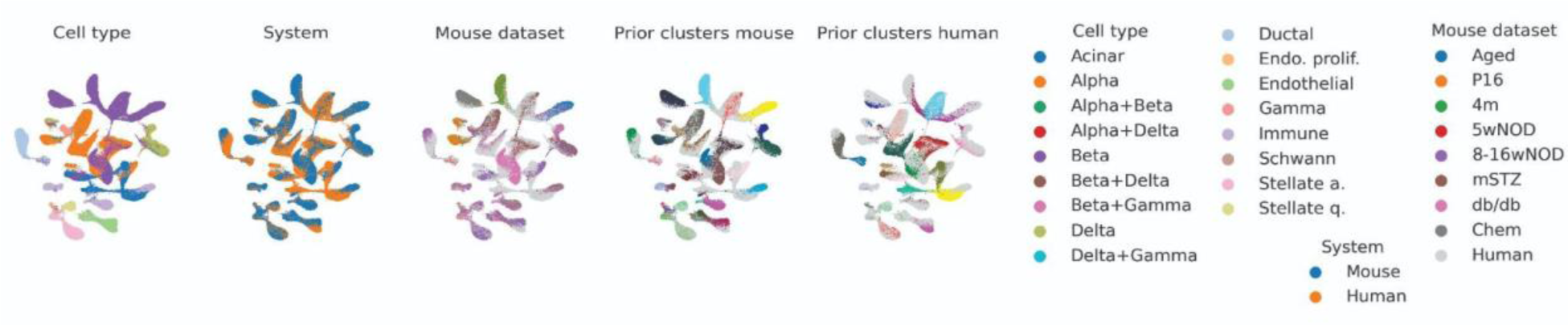
Batch effects in prior cluster labels are reflected in the final SATURN integration. The UMAPs display the per-system prior clusters and the dataset covariate for the mouse system. They were computed on a representative run from the best hyperparameter setting for SATUR mouse-human integration. Prior cluster color legend is not shown as individual cluster names are not meaningful. For the mouse-specific covariates (datasets and clusters) the human data is shown as a background in light gray, and vice versa for the human covariates.

**Supplementary Figure S15:**
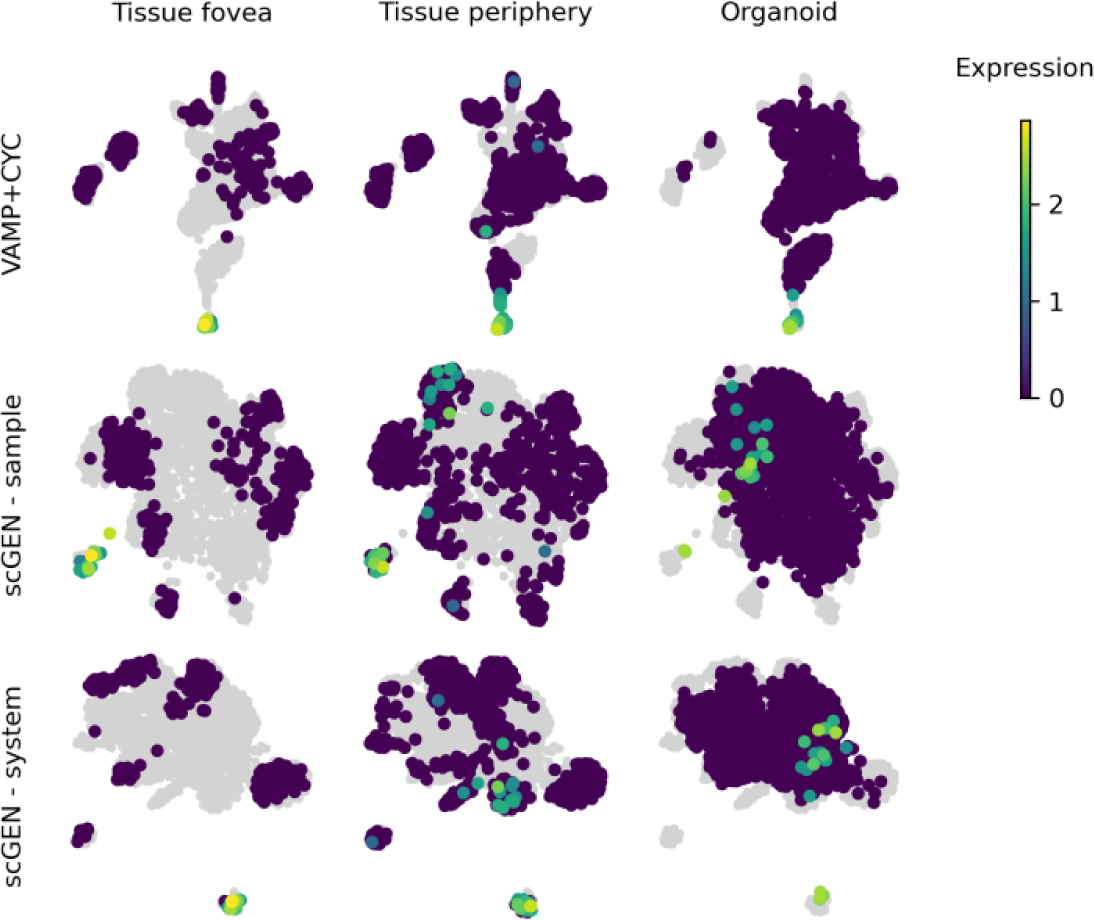
Cell subtype preservation in amacrine cells after integration. The UMAPs are colored by expression of starburst amacrine cell marker *SLC18A3*. They were computed for the amacrine cell subset from the organoid-tissue dataset based on a representative run for the best hyperparameter setting of VAMP+CYC and on two example runs of scGEN with samples or systems as batch covariates.

**Supplementary Figure S16:**
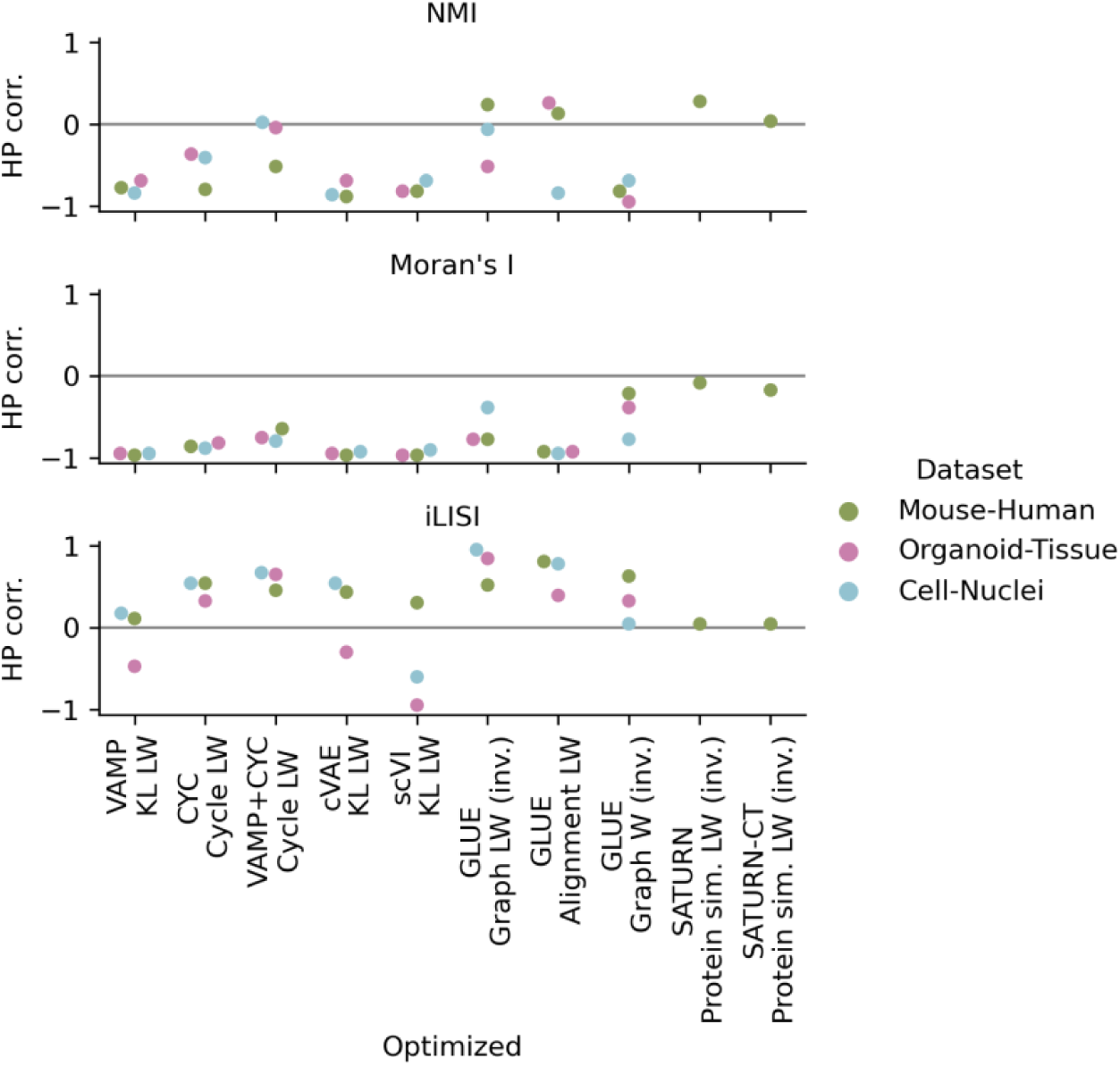
Spearman correlation between hyperparameter values and integration metrics. Correlation (corr.) was computed for different combinations of models and tuned hyperparameters (x-axis) across all runs and hyperparameter values (HP) in the OTO setting as shown in **Supplementary Figure S1**, separately for each dataset. Values of hyperparameters that are expected to be negatively rather than positively associated with batch correction strength were inverted (inv.) before computing correlation.

## Supplementary tables

**Supplementary Table S1: Comparison of pre-integration sample distances within and between systems.** One-sided Mann–Whitney U test was used to compare distributions of per-cell-type sample distances in the PCA space within a system (control) and between systems (case) for every data use cases (sheet names). Column names: cell_type - tested cell type; system - control group, with “within/between” in the mouse-human data referring to within/between dataset distances in the mouse data; u - test statistic; padj - adjusted p-value; n_system and n_crossystem - number of distances (observations) in the control and case groups, respectively.

**Supplementary Table S2: Statistical comparison of model performance in different data use cases.** Welch’s t-test was used to compare integration performance between different models in individual data use cases. For each model, the best hyperparameter setting was selected as described in the methods and shown in Figure 5. Column names: dataset - the integrated dataset, metric - integration metric, model - the used integration model, p - p-value, t - test statistic, padj_fdr_tsbh - adjusted p-value, model_cond - model used as the test condition, model_ctrl - model used as the control condition, higher - model with higher value of the tested metric.

